# SCD-dependent lipid metabolism licenses alternative macrophage activation and macrophage plasticity

**DOI:** 10.64898/2026.05.27.728175

**Authors:** Tianheng Hou, Richard L. Watson, Alexander H. Bedard, Wei-Yuan Hsieh, Jonathan Quintana, Baolong Su, Amara C. Thind, Juliette N. Uy, Clay F. Semenkovich, Peter J. Bradley, Peter Tontonoz, Kevin J. Williams, Autumn G. York, Steven J. Bensinger

**Affiliations:** Department of Microbiology, Immunology, and Molecular Genetics, University of California, Los Angeles, Los Angeles, CA 90095, USA; Division of Pulmonary & Critical Care Medicine, Department of Medicine, University of California Los Angeles, CA 90095, USA; Section of Pulmonary & Critical Care, Department of Medicine, VA Greater Los Angeles Healthcare System, Los Angeles, CA 90073, USA; Department of Pathology and Laboratory Medicine, University of California, Los Angeles, Los Angeles, CA 90095, USA; Department of Biological Chemistry, University of California, Los Angeles, Los Angeles, CA 90095, USA; UCLA Lipidomics Laboratory, University of California, Los Angeles, Los Angeles, CA 90095, USA; Division of Endocrinology, Metabolism and Lipid Research, Washington University School of Medicine, St. Louis, MO 63110, USA; Department of Immunology, School of Medicine, University of Washington, Seattle, WA 98195, USA; Department of Molecular and Medical Pharmacology, David Geffen School of Medicine, University of California, Los Angeles, CA 90095, USA; Department of Immunology and Immune Therapeutics, Keck School of Medicine, University of Southern California, Los Angeles, CA 90033, USA

**Keywords:** macrophage, alternative activation, lipid metabolism, macrophage plasticity, toxoplasma

## Abstract

Lipid metabolic reprogramming accompanies macrophage activation, yet our understanding of why macrophages profoundly reshape their lipid composition remains unclear. Here, we identify stearoyl-CoA desaturase (SCD) as a lipid-metabolic checkpoint required for the acquisition of the alternatively activated macrophage (AAM) cell state. We show that SCD maintains lipid desaturation balance by ensuring the conversion of newly synthesized saturated long-fatty acids (SFAs) into monounsaturated fatty acids (MUFAs). Critically, disruption of SCD resulted in the rapid accumulation of saturated phosphatidic acid (PA) and hyperactivation of mTORC1, which perturbed the acquisition of stable AAM epigenetic and transcriptional programs. Notably, inhibition of upstream lipogenic enzymes (e.g., ACC and FASN) did not impair AAM polarization, underscoring that an imbalance between newly synthesized saturated and monounsaturated fatty acids, not the loss of *de novo* lipogenesis, determines AAM identity. Unexpectedly, the loss of SCD redirected IL-4-activated macrophages into an aberrant cell state poised for pro-inflammatory responses, indicating that disruption of this lipid metabolic checkpoint leads to misinterpretation of anti-inflammatory instructional cues. Rescue of SCD-deficient macrophages via mTORC1 inhibition confirmed a causal role for the SCD-PA-mTORC1 axis in controlling macrophage plasticity. Finally, we find that SCD deficiency and resultant PA accumulation attenuated *Toxoplasma gondii*-driven AAM polarization and reduced pathogen burden. Together, these data establish the SCD-PA lipid signaling axis as a critical metabolic regulator of macrophage plasticity with direct consequences for host defense.

## INTRODUCTION

Macrophages are specialized immune cells that generate inflammatory responses, eliminate harmful stimuli, defend the host from pathogens, clear extracellular debris and dead cells, and coordinate tissue repair[1–3]. Consequently, macrophages are highly plastic cells, that continuously integrate immune, environmental, and metabolic cues to acquire specific cellular states supporting distinct effector programs. This phenomenon, described initially as polarization into M1 or M2 states in response to classical pro- or anti-inflammatory stimuli (e.g., LPS or IL-4)[4, 5], is now recognized as a dynamic continuum of activation states determined by the nature of the instructive signals received. Macrophages can also transition between discrete cell states over time as they receive and integrate new information. For example, macrophages initially adopting a pro-inflammatory phenotype in response to TLR activation may later shift towards a wound healing or pro-resolving state in response to immunoregulatory cytokines (e.g., IL-10, TGF-β). Such plasticity is essential for limiting inflammation and promoting resolution. There is also evidence that initial treatment with immunoregulatory type II cytokines (e.g., IL-4) can prime macrophages for heightened pro-inflammatory responses when subsequently activated with TLR signals[6]. Thus, macrophage plasticity is an intrinsic property that ensures proper function. Consequently, restricting the movement of macrophages between distinct cellular states contributes to pathological inflammatory responses and disease[7, 8].

Macrophage polarization is tightly linked with changes in cellular lipid metabolism. Upon activation, macrophages reshape their lipid composition, or lipidomes, in a stimulus-specific manner[9]. It is thought that changes in lipid composition are necessary to support effector programs, and accordingly, restricting the ability of macrophages to acquire their “preferred” lipidomes can influence their function and drive disease pathogenesis[10]. The long-chain fatty acid biosynthetic pathway, or *de novo* lipogenesis (DNL), has been implicated in regulating polarization into an alternatively activated macrophage (AAM) state[11]. IL-4 and IL-10 signaling upregulates DNL, and the loss of SREBP1 and PPARγ, key transcriptional regulators of DNL, can attenuate AAM polarization[12, 13]. However, studies that targeted specific enzymes of the DNL pathway yielded conflicting results. Genetic deletion of acetyl coenzyme A carboxylase 1 (ACC1), the enzyme responsible for synthesis of malonyl coenzyme A in long-chain fatty acid biosynthesis, did not influence IL-4-mediated AAM polarization[14], whereas loss of fatty acid synthase (FASN), the rate-limiting enzyme of long-chain fatty acid biosynthesis, impaired AAM polarization, though this effect was attributed to alterations in redox state rather than specific changes in lipid composition[11]. Thus, the question of whether DNL-mediated reshaping of macrophage lipid composition is required for AAM polarization and effector function remains indeterminate.

To address this gap in our knowledge, we applied multi-omic approaches and identified the 19-stearoyl-CoA desaturases (SCD1 and SCD2 in mouse, and SCD1 in humans), enzymes responsible for the synthesis of long-chain monounsaturated fatty acids (MUFAs), as essential for AAM polarization in mouse and human macrophages. In the absence of SCDs, IL-4-activated macrophages failed to fully acquire an AAM program, and instead, were diverted into an aberrant cell state poised for pro-inflammatory responses. Notably, perturbation of upstream DNL enzymes (e.g., ACC1 or FASN) had minimal effects on AAM polarization, indicating that the balance between saturated and monounsaturated fatty acid synthesis, rather than total lipogenic flux, is central to acquiring AAM identity. Mechanistically, impaired AAM polarization in SCD-deficient macrophages was linked to the abnormal accumulation of saturated phosphatidic acids, which led to sustained mTORC1 activation. Finally, we demonstrate that disrupting SCD, and the resulting accumulation of saturated PAs, impaired the pathogen-driven conversion of macrophages into an alternatively activated state during *Toxoplasma gondii (T. gondii)* infection. Together, these findings identify the SCD–PA lipid metabolic axis as a critical regulator of macrophage plasticity and provide proof-of-concept that targeting this metabolic axis can render macrophages refractory to pathogen-directed changes in functional state.

## RESULTS

### Perturbing monounsaturated fatty acid biosynthesis disrupts AAM polarization

The SCD enzymes convert synthesized or imported long-chain saturated fatty acids (SFAs) into monounsaturated fatty acids (MUFAs)[15]. We previously reported that IL-10 simulation of pro-inflammatory macrophages enhanced SCD2-mediated MUFA synthesis, which was required to attenuate inflammation[16]. These observations led us to ask the broader question of whether reprogramming MUFA synthesis is essential for alternative macrophage polarization. Murine cells express multiple SCD isoforms (SCD1-4)[15]. To identify the relevant SCD enzymes, IL-4-activated bone marrow derived macrophages (BMDMs) from C57BL/6 (WT) mice were cultured in complete media and gene expression assessed. We found that that *Scd1* and *Scd2* were upregulated during AAM polarization, while *Scd3* and *Scd4* were not expressed at appreciable levels (Fig, S1a). Thus, we used co-deletion of *Scd1* and *Scd2* in our murine genetic studies hereafter.

Initially, we tested whether co-deletion of *Scd1* and *Scd2* in macrophages perturbed MUFA homeostasis during AAM polarization. To that end, *LysMCre* (LysM-control) and *LysMCre-SCD1/2^fl/fl^* BMDMs (designated SCD1/2 KO) were stimulated with IL-4 for 48 h in complete media containing 50% 13C-glucose. Lipids were extracted, derivatized, and GC/MS performed to assess ^13^C-isotopic enrichment into newly synthesized SFAs (e.g., 16:0) and MUFA (e.g.,18:1) synthesized during polarization. We found that IL-4 activation increased the amount of palmitate and oleate synthesized (Fig. S1b). As expected, genetic deletion of *Scd1* and *Scd2* markedly decreased MUFA synthesis, resulting in a shift in the desaturation index (ratio of 18:1 to 18:0) of IL-4-activated macrophages (Fig. S1b). Similarly, treating WT BMDMs with a small molecular inhibitor of SCD (Cay10566; 2-10nM; designated SCDi hereafter) also attenuated the upregulation of MUFA synthesis and shifted the desaturation index of IL-4-activated macrophages (Fig.S1c).

Next, we asked whether genetic or pharmacologic disruption of MUFA synthesis perturbed AAM polarization. To initially test this question, LysM-control and SCD1/2 KO BMDMs were polarized with IL-4 as above. After 48 h, macrophages were collected and assessed by flow cytometry (FACS) and qPCR to determine polarization into AAMs. As expected, IL-4-activated LysM-control macrophages upregulated expression of canonical AAM-associated proteins CD301, RELMα, and Arginase 1(Arg1), with ∼ 40% of the cultures becoming CD301+/RELMα+ or CD301+/Arg1+ (Fig. 1a,b and S1d,e). Gene expression confirmed robust upregulation of hallmark AAM genes *Arg1*, *Ym1*, *Mgl2*, and *Retnla* (Fig. 1c). Remarkably, SCD1/2 KO macrophages did not efficiently polarize into AAMs as evidenced by a 4-fold decrease in the frequency of CD301+/ RELMα+ or CD301+/Arg1+ macrophages (Fig. 1a,b and S1d,e), and markedly attenuated IL-4-induced upregulation of *Arg1*, *Ym1*, *Mgl2*, and *Retnla* (Fig. 1c). We considered the possibility that genetic loss of SCD1/2 predisposed macrophages to be refractory to IL-4-mediated polarization. To address this, we pharmacologically inhibited SCD (Cayman 10566 designated SCDi; 10nM)[9] to acutely perturb this lipid metabolic pathway in BMDMs. We observed a similar impact on AAM polarization at the phenotypic and gene expression level in WT IL-4-activated macrophage cultures (Fig. 1d,e,f and S1f,g). Importantly, providing SCD1/2-deficient or inhibited macrophage cultures with exogenous oleate (18:1), the primary product of SCD, restored AAM polarization (Fig. 1a-f and S1d-g). Together, these data show a critical role for MUFA synthesis in AAM polarization.

**Figure 1.**
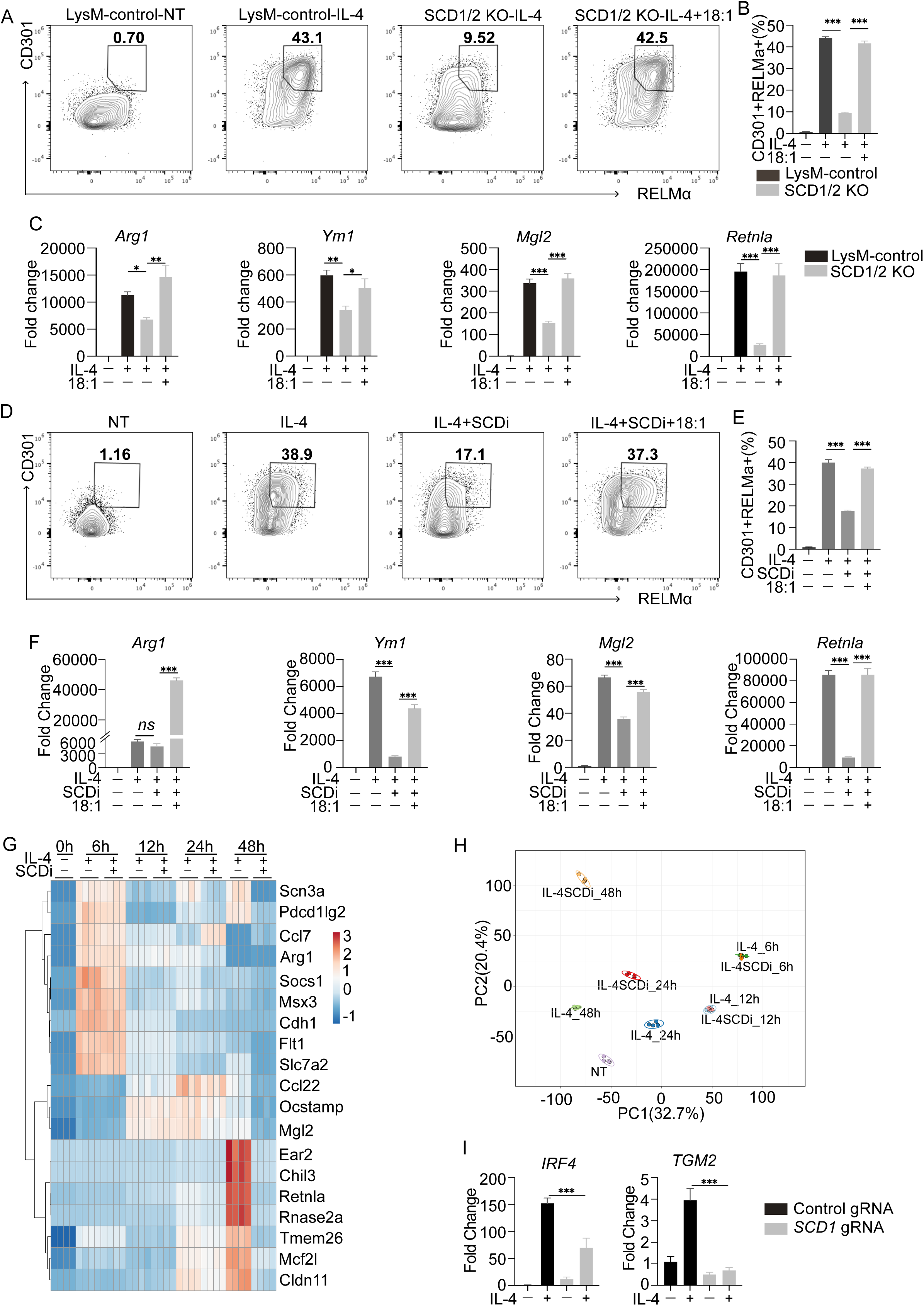
Loss of MUFA synthesis by the stearoyl-CoA 1 and 2 (SCD1/2) enzymes disrupt AAM polarization. A. Flow cytometry plots of LysMCre control stimulated with or without IL-4 and LysMCre-SCD1/2^fl/fl^ BMDMs stimulated with IL-4 ± BSA-conjugated 18:1 fatty acid (hereafter as 18:1, 25 μM) as indicated for 48 h. B. Bar graph depicts percent of double positive (CDCD301+/RELMα+) BMDMs in A. C. qPCR analysis of indicated AAM-associated genes from LysMCre control with or without IL-4 and LysMCre-SCD1/2^fl/fl^ BMDMs stimulated with IL-4 ± 18:1 (25 μM) as indicated for 48 h. D. Flow cytometry plots of quiescent BMDMs (NT) and BMDMs stimulated with IL-4 or IL-4+SCDi (Cay10566) ± 18:1(25 μM) as indicated for 48 h. E. Bar graph depicts percent of double positive (CDCD301+/RELMα+) BMDMs in D. F. qPCR analysis of indicated AAM-associated genes from BMDMs (NT) and BMDMs stimulated with IL-4 or IL-4+SCDi (Cay10566) ± 18:1(25 μM) as indicated for 48 h. G. Heatmap of selected AAM associated genes from RNA-seq data of quiescent BMDMs (NT) or BMDMs stimulated with IL-4 ± SCDi for 0h, 6h, 12h, 24 h and 48h. Genes (rows) were clustered according to correlation. H. Principal component analysis (PCA) of RNA-seq data from the same conditions in G. Percentage of total variance explained by individual principal components (PC1 and PC2) indicated in axis. Prediction ellipses are set at 95% probability (n = 4 replicates/experimental condition). I. qPCR analysis of indicated AAM-associated genes from human monocyte-derived macrophages with control or CRISPR-mediated SCD1 deletion, stimulated with or without human IL-4 for 48 h. Gene expression and flow cytometry studies are from three biologic replicates per experimental condition and are representative of greater than two experiments. All bar graphs are presented as mean ± SEM. *p < 0.05; **p < 0.01, ***p < 0.001.

To better understand how loss of MUFA synthesis altered the AAM gene expression program, WT macrophages were stimulated with IL-4 alone or in combination with SCDi (10nM), and transcriptomics was performed on T = 0 h (designated NT), 6 h, 12 h, 24 h, and 48 h samples. Analysis of a panel of AAM-associated genes showed that SCD inhibition did not alter expression of these AAM genes at early time points (6 h, 12 h, 24 h) (Fig. 1g). In contrast, the upregulation of AAM genes at later time points (e.g., *Ear2*, *Chil3*, *Retnla*) was markedly attenuated by SCD inhibition. We also assessed how loss of MUFA synthesis changes the global gene expression program of IL-4-stimulated macrophages. Clustering and principal component analysis (PCA) showed that SCD inhibition had minimal influence on the transcriptome of IL-4-activated macrophages at 6 h and 12 h (Fig. 1h and S1h). However, a divergence in the gene expression program of SCD-inhibited macrophages was apparent at the 24 h time point, and this difference widened further at 48 h (Fig. 1h and S1h). Pathway analysis of differentially expressed genes (DEGs) at 24 h revealed the emergence of an NF-κB and inflammatory gene signature, along with a cellular stress response (Fig. S1i). Few downregulated genes were observed and were primarily related to changes in fatty acid homeostasis (Fig. S1i). Analysis of DEGs at 48 h showed a continued inflammatory gene signature, and the upregulation of the ER stress program (Fig. S1j). Notably, AAM-associated genes were amongst the most significantly down-regulated genes at this latter time point. We also performed CRISPR-mediated reduction in SCD1 expression in human monocyte-derived macrophages (Fig. S1k). We found that loss of SCD1 rendered human macrophages resistant to AAM polarization, as evidenced by attenuated upregulation of *IRF4* and *TGM2* in response to IL-4 activation (Fig. 1i). Thus, loss of SCD enzymes, and the corresponding loss of MUFA synthesis, perturbs the AAM polarization potential of mouse and human macrophages.

We considered the possibility that loss of SCD function perturbed IL-4R/STAT6 signaling in macrophages. However, we found that neither genetic deletion of SCD nor pharmacologic inhibition of SCD attenuated STAT6 phosphorylation (Fig. S1l,m). Therefore, perturbed IL-4R/STAT6 signaling cannot account for the decrease in AAM polarization potential observed when SCD-mediated MUFA synthesis is perturbed. Together, these data show that MUFA biosynthesis via SCD enzymes is essential for polarization of AAMs, and that loss of MUFA biosynthesis rewires the transcriptional response of IL-4 activated macrophages.

### Loss of SCD redirects IL-4 polarized macrophages into an aberrant cell state

Our data indicate that genetic or pharmacologic inhibition of SCD alters the transcriptional program of IL-4-activated macrophages by 24 h after stimulation. Thus, we asked if loss of MUFA biosynthesis influenced the epigenetic state. To address this, BMDMs were stimulated with IL-4 with or without SCDi for 24 h, followed by assay of transposase accessible chromatin sequencing (ATAC-seq). A global analysis of transcription start sites (TSS) did not reveal differences in general TSS accessibility across experimental conditions (Fig. 2a). Next, we applied a sliding window approach to identify differentially accessible regions (DARs) among the experimental conditions (Fig. S2a,b) and performed motif analysis[17]. We initially focused on the differences between unstimulated and IL-4-stimulated macrophages to identify the transcription factor binding motifs that were most accessible as a consequence of AAM polarization (Fig. 2b and Supp Table 1)[18, 19]. As expected, among the most significantly accessible motifs, we observed opening of chromatin at binding sites for STAT6, a master transcriptional regulator of alternative macrophage activation (Fig. 2b, Supp Table 1)[18]. Similarly, we found increased accessibility of early growth response (EGR) 1 and 2 motifs, consistent with previous studies showing that these TFs are upregulated by IL-4 or IL-13 and play a critical role in the polarization and function of AAMs[20]. Our analysis also revealed increased chromatin accessibility at TF binding sites associated with regulation of macrophage activation and polarization, such as the activator protein 1 (AP-1) family members (e.g., JUN, FOS, BATF)[21, 22]. Increased accessibility of Activating Transcription Factor 3 (ATF3) binding sites was also noted, which has also been shown to promote pro-resolving or anti-inflammatory macrophage states[23].

**Figure 2.**
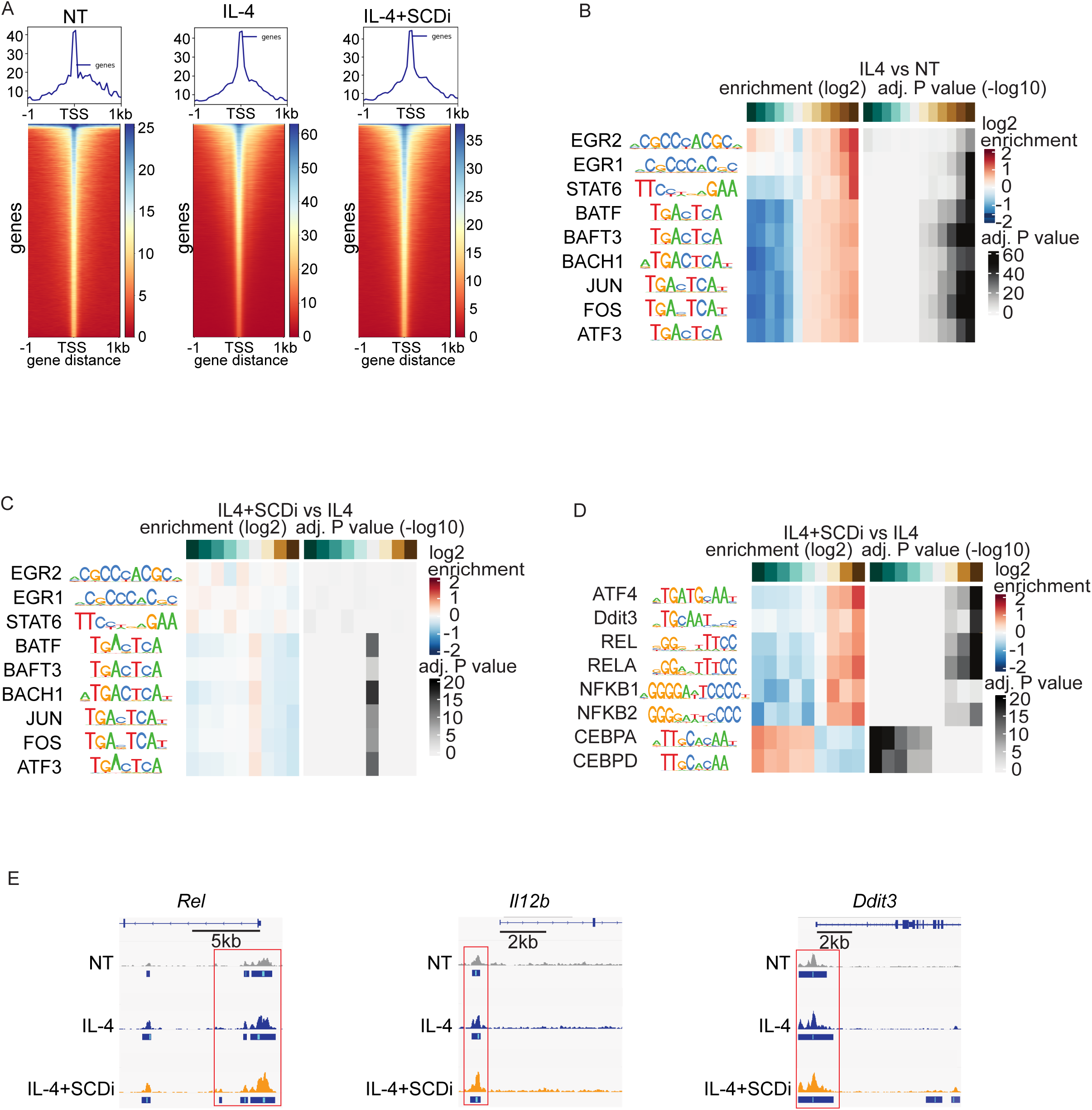
SCD1/2 is required for canonical AAM chromatin remodeling and loss of SCD1/2 results in an aberrant cell state. A. Representative plot and heatmap analysis of ATAC-seq signal within ±1kb of the transcription start site (TSS) in quiescent BMDMs (NT) or BMDMs stimulated with IL-4 ± SCDi for 24 h. B. Comparison of motif binding sites identified in quiescent BMDMs (NT) or BMDMs stimulated with IL-4 for 24 h. Heatmaps representing enrichment (left) and adjusted p-values (right) for each of the sliding window bins (top). C. Comparison of AAM associated motifs in BMDMs stimulated with IL-4 ± SCDi for 24 h. Heatmaps representing enrichment (left) and adjusted p-values (right) for each of the sliding window bins (top). D. Comparison of motif binding sites identified in BMDMs stimulated with IL-4 ± SCDi for 24 h. Heatmaps representing motif enrichment (left) and adjusted p-values (right) for each of the sliding window bins (top). E. Representative IGV browser tracks displaying normalized ATAC-seq peak counts at *Rel*, *Il12b,* and *Ddit3* loci in quiescent BMDMs (NT) and BMDMs stimulated with IL-4 ± SCDi for 24 h. ATAC-seq experiments are combined from three biologic replicates per experimental condition. Tracings are representative.

Having defined a core set of AAM motifs regulated by IL-4 in our model system, we then assessed whether inhibiting SCD during IL-4 polarization altered the chromatin accessibility at these sites. We found that SCD inhibition resulted in a modest, but not significant, closing of STAT6 motif sites (Fig 2c and Supp Table 1). No difference in accessibility of the other IL-4 responsive binding sites (e.g., EGR, JUN, FOS, BATF, and ATF3) was noted. We also considered the possibility that loss of SCD was constraining chromatin accessibility for other TF reported as required for AAM polarization, such as IRF4, PPARγ, KLF-4, MafB, and c-Myc, however, loss of SCD also did not alter chromatin accessibility of these TF motif sites (Supp Table 1). Thus, the loss of SCD-mediated MUFA synthesis does not appear to grossly impact chromatin accessibility for canonical AAM-associated TF binding motifs at 24 h (Fig 1g).

Next, we determined how loss of MUFA synthesis more broadly influenced chromatin accessibility during AAM polarization. Analysis revealed that SCD inhibition markedly increased accessibility for binding sites of multiple NF-κB family members (e.g., REL, RELA, NFκB1, NFκB2) and ER stress factors (e.g., ATF4, Ddit3) (Fig. 2d). These changes are consistent with our pathway analysis of RNA-Seq data at this same time point, which indicated that inhibition of SCD activity increases the expression of pro-inflammatory and ER stress genes (Fig. S1j). In contrast, analysis revealed that binding motifs for the CCAAT-enhancer-binding protein (C/EBP) family were the most significantly constrained (Fig. 2d, Supp Table 1). Of potential importance, C/EBP proteins have been shown to play critical roles as transcriptional regulators in macrophage identity and specifying the AAM gene program[24]. Examination of select genomic loci confirmed that SCD inhibition was associated with increased accessibility of *Rel*, NF-κB target gene *Il12b*, and ER stress response gene *Ddit3* when compared to IL4 stimulation alone (Fig. 2e).

Finally, we noted that SCD inhibition increased accessibility of chromatin regions with higher GC-content compared to IL-4 treatment alone (Supp Fig2c). Given that GC-enrichment and CpG content can potentially bias the assessment of motif enrichment, we performed stability selection (LASSO regularization) to account for this possibility[25]. Similar to our initial motif analysis (Fig 2d), we observed that NF-κB (REL, RELA) and ER stress (ATF4, Ddit3) emerged as the most significantly enriched binding sites when SCD was inhibited (Fig. S 2d). Furthermore, loss of SCD resulted in closure of CEBP-A/D binding sites (Fig. S 2d). In combination with our transcriptomics studies, our data indicate that inhibition of SCD during IL-4 polarization redirects macrophages into an altered, pro-inflammatory cell state, incapable of effectively supporting an AAM phenotype.

### Inhibiting saturated long chain fatty acid synthesis does not perturb AAM polarization

Saturated LCFAs (e.g., palmitate and stearate) are the substrate of SCD enzymes and can be imported or synthesized *de novo* (Fig. 3a). Thus, we asked if disrupting saturated LCFA biosynthesis would phenocopy our results with SCD inhibition and attenuate AAM polarization. To address this, *LysM-*control and *LysMCre-Fasn^fl/fl^* BMDMs (FASN KO) were activated with IL-4 as above for 48 h and then assessed for hallmark characteristics of AAMs. Flow cytometry studies of macrophages revealed no difference in the frequency of AAM-associated CD301+/ RELMα+, CD301/Arg1+ or CD206+/71+ macrophages (Fig. 3b,c and S3a). Likewise, gene expression studies for hallmark AAM genes showed that FASN KO macrophage were able to upregulate *Arg1*, *Ym1*, *Mgl2*, and *Retnla* similar to control macrophages (Fig. 3d). We were surprised by this result and considered the possibility that *LysMCre-Fasn^fl/fl^* BMDMs retained expression of FASN in our studies. Consistent with this, immunoblots revealed FASN KO BMDMs retained ∼50% of FASN protein relative to controls (Fig. 3e), corresponding to a similar (∼50%) decrease in palmitate (16:0) synthesis (Fig. S3b). Thus, we conclude that conditional deletion of *Fasn* using *LysMCre* was not efficient enough in our model to rigorously test the importance of saturated LCFA biosynthesis in AAM polarization.

**Figure 3.**
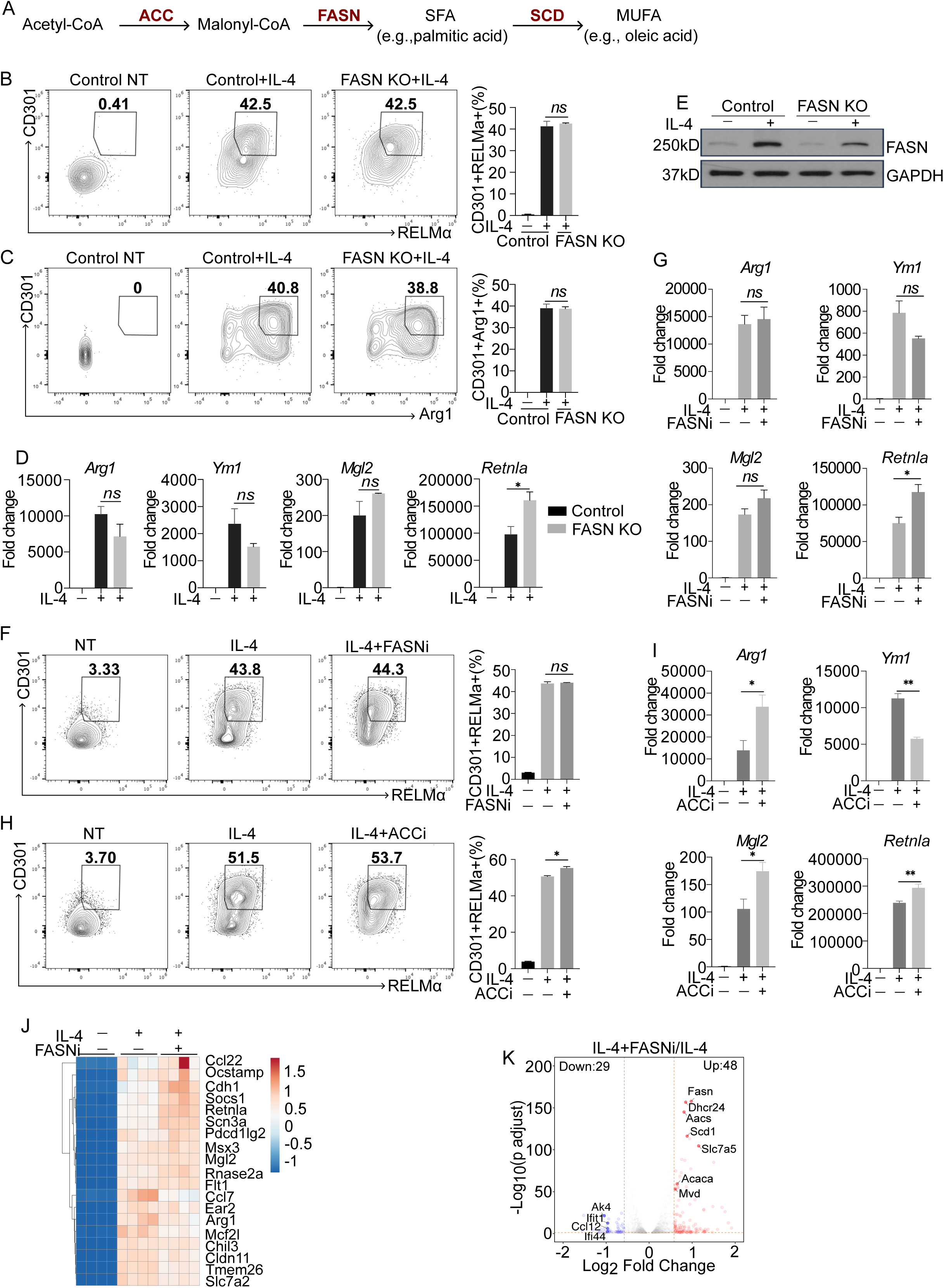
Fatty acid synthetic enzymes ACC and FASN are not required for AAM polarization in vitro. A. Simplified schematic of de novo fatty acid synthesis pathway. B. Flow cytometry plots of LysMCre control stimulated with or without IL-4 and LysMCre-FASN^fl/fl^ BMDMs stimulated with IL-4 as indicated for 48 h. Bar graph (right) depicts percent of double positive (CDCD301+/RELMα+) BMDMs. C. Flow cytometry plots of LysMCre control stimulated with or without IL-4 and LysMCre-FASN^fl/fl^ BMDMs stimulated with IL-4 as indicated for 48 h. Bar graph (right) depicts percent of double positive (CDCD301+/Arg1+) BMDMs. D. qPCR analysis of indicated AAM-associated genes from LysMCre control stimulated with or without IL-4 and LysMCre-FASN^fl/fl^ BMDMs stimulated with IL-4 as indicated for 48 h. E. Western blot analysis of FASN from LysMCre-control (Control) and LysMCre-Fasn^fl/fl^ BMDMs stimulated with or without IL-4 for 48 h. F. Flow cytometry plots of quiescent BMDMs (NT) and BMDMs stimulated with IL-4 ± FASN inhibitor (TVB-3664; 10nM; indicated as FASNi) as indicated for 48 h. Bar graph (right) depicts percent of double positive (CDCD301+/RELMα+) BMDMs. G. qPCR analysis of indicated AAM-associated genes of quiescent BMDMs (NT) and BMDMs stimulated with IL-4 ± FASNi inhibitor as indicated for 48 h. H. Flow cytometry plots of quiescent BMDMs (NT) and BMDMs stimulated with IL-4 ± ACC inhibitor (ND-646; 100nM; indicated as ACCi) as indicated for 48 h. Bar graph (right) depicts percent of double positive (CDCD301+/RELMα+) BMDMs. I. qPCR analysis of indicated AAM-associated genes of quiescent BMDMs (NT) and BMDMs stimulated with IL-4 ± ACCi inhibitor as indicated for 48 h. J. Heatmap of selected AAM associated genes from RNA-seq data of quiescent BMDMs (NT) or BMDMs stimulated with IL-4 ± FASNi for 24 h. Genes (rows) were clustered according to correlation. K. Volcano plot of differentially expressed genes (DEGs) from RNA-seq analysis of BMDMs stimulated with IL-4 ± FASNi for 24 h. All experiments are from four biologic replicates per experimental condition. Heatmap scales are z-score for number of deviations away from the row mean. Rows are clustered using correlation distance and complete linkage. Gene expression and Flow cytometry studies are from three biologic replicates per experimental condition and are representative of greater than three experiments. Bar graphs are presented as mean ± SEM. *p < 0.05; **p < 0.01, ***p < 0.001.

To overcome this barrier, we treated WT BMDMs with small molecule inhibitors of FASN (e.g., TVB-3664 or C75). Dose-response studies using ^13^C-stable isotope enrichment into palmitate (16:0) were performed (as above in Figs. S1b and S1e) to ensure that inhibition of saturated LCFA synthesis was achieved using the lowest dose possible. The IC95 (95% inhibitory concentration) for palmitate (16:0) synthesis by TVB-3664 was calculated to be 9.2 nM (Fig. S3c). Unexpectedly, treating macrophages with the FASN inhibitor C75 did not inhibit palmitate synthesis at the doses tested (0.1-50 mM) under these polarization conditions (Fig. S3d). For subsequent studies, we used TVB-3664 at 10nM (designated FASNi hereafter). WT macrophages were polarized with IL-4 alone or in combination with TVB-3664 (10nM) and assessed by flow cytometry for acquisition of the AAM phenotype. FACS analysis revealed no difference in the frequency of CD301+/RELMα+, CD301/Arg1+ or CD206+/71+ macrophages between control and FASNi treatment (Fig. 3f, S3e, f). Similarly, FASN inhibition did not interfere with upregulation of AAM genes *Arg1*, *Ym1*, *Mgl2 and Retnla* (Fig. 3g).

We found this result surprising and tested whether inhibition of ACC1, the enzyme responsible for generating malonyl-CoA used by FASN in the production of saturated LCFA, would perturb AAM polarization. Does response curves with the ACC1 inhibitor ND646 (designated ACCi hereafter) indicated 80-100nM completely blocked palmitate synthesis in IL-4 activated macrophages (Fig. S3g). Similar to our studies with FASN inhibited macrophages, ACC1i treatment at 100nM did not alter the frequency of CD301+/RELMα+, CD301/Arg1+, or CD206+/71+ macrophages (Fig. 3h and S3h,i). Moreover, gene expression studies suggested ACC1 inhibition promoted the expression of *Arg1*, *Mgl2 and Retnla* (Fig. 3i), however *Ym1* was decreased (Fig. 3i). Together these data indicate that disrupting DNL does not grossly influence AAM polarization.

As a cautionary note, we found that treating macrophages with the FASN inhibitor C75 attenuated upregulation of hallmark AAM genes (e.g., *Arg1*, *Mgl2, Ym1*, *Retnla*) in IL-4-polarized macrophages in a dose-dependent manner (at 2.5-10 μM Fig. S3j), in keeping with prior reports[11]. Thus, we conclude that C75’s inhibitory influence on AAM polarization is independent of LCFA synthesis and is likely an off-target effect.

To gain a better understanding of how inhibition of saturated LCFA synthesis altered the gene expression program and epigenetic state during AAM polarization, we performed bulk RNA-seq and ATAC-seq. In these studies, WT macrophages were activated with IL-4 alone or in combination with FASNi (TVB-3664; 10nM) for 24 h before collection. Consistent with flow cytometry and qPCR studies, inhibition of FASN did not influence the expression of the AAM-associated genes (Fig. 3j). Inspection of the broader transcriptional program indicated that disrupting FASN during IL-4 polarization had minimal influence on the AAM gene expression program, with only 77 genes (48 up and 29 down) reaching statistical significance relative to IL-4 alone (Fig. 3k, Fig S3k). Inspection of genes upregulated in response to FASN inhibition showed enrichment for lipid biosynthesis (e.g., *Fasn, Scd1, Aacs, Dhcr24*), suggesting a compensatory increase in lipid metabolic processes (Fig. 3k). Analysis of ATAC-seq data showed no statistical difference in chromatin accessibility at TF binding sites associated with AAM polarization (e.g., STAT6, PPARγ) (Fig. S3l) nor did we see changes in chromatin accessibility driven by SCD inhibition (e.g., ATF4, RELA, CEBP) (Fig S3l). Together, these data indicate that saturated LCFA synthesis is dispensable for the acquisition of the AAM cell state with the polarizing conditions used here.

### Disruption of MUFA synthesis drives the accumulation of phosphatidic acid (PA) thereby perturbing AAM polarization

Next, we focused on elucidating the molecular mechanism by which disrupting MUFA synthesis limits macrophage polarization into AAMs. Our data indicated that replenishing oleate (18:1) in cultures was sufficient to restore AAM polarization in SCD-loss-of-function models (Fig. 1). Thus, we hypothesized that the lipid(s) responsible for interfering with AAM polarization when SCD-mediated MUFA synthesis was lost, would be normalized when oleate was supplemented to cultures. To test this hypothesis, we performed lipidomics on IL-4-activated *LysM*-control and SCD1/2 KO macrophages alone or with supplemental oleate. After 48 h, lipidomics was performed to assess changes in lipid composition. Cluster analysis revealed a distinct group of lipids that accumulated in SCD1/2 KO macrophages during AAM polarization and were normalized by oleate supplementation (Figure 4a; blue box). Inspection of these lipids showed enrichment for phosphatidic acid (PA), diacylglycerols (DAG), lysophosphatidylcholines (LPC), and hexosyl ceramides (HexCer) species. Analysis of total amounts of these lipid subclasses showed a similar pattern, with accumulation of total PA showing the largest change (Fig. 4b, S4a). PA32:0 and PA34:0 accounted for nearly 80% of the total PA accumulated (Fig. 4c).

**Figure 4.**
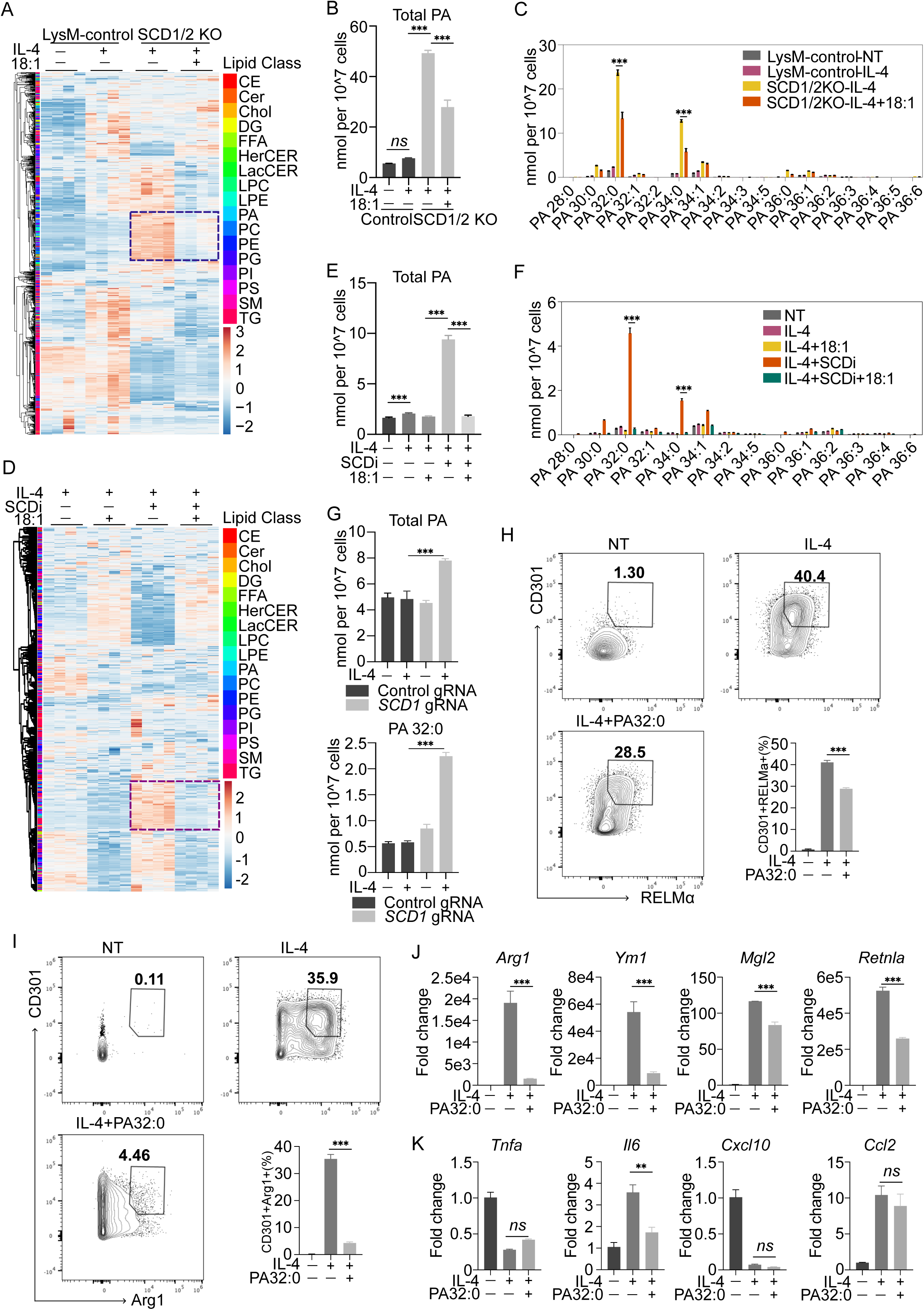
PA homeostasis is essential for the acquisition of AAM. A. Heat map of individual lipids quantified by shotgun lipidomics from LysMCre control BMDMs stimulated with or without IL-4 and LysMCre-SCD1/2^fl/fl^ BMDMs stimulated with IL-4 ± 18:1 (25 μM) as indicated for 48 h. B. Total phosphatidic acid (PA) measured by shotgun lipidomics from LysMCre control BMDMs stimulated with or without IL-4 and LysMCre-SCD1/2^fl/fl^ BMDMs stimulated with IL-4 ± 18:1 (25 μM) as indicated for 48 h. C. Amounts of individual PA species measured by shotgun lipidomics from BMDMs stimulated as above. D. Heat map of individual lipids quantified by shotgun lipidomics from BMDMs stimulated with IL-4 ± 18:1 (25 μM) or IL4 with SCDi ± 18:1 (25 μM) as indicated for 48 h. E. Total phosphatidic acid (PA) measured by shotgun lipidomics from quiescent BMDMs (NT) or BMDMs stimulated with IL-4 ± SCDi or IL-4 ± SCDi with 18:1 (25 μM) as indicated for 48 h. F. Amounts of individual PA species measured by shotgun lipidomics from BMDMs stimulated as above. G. Total phosphatidic acid (PA) (up) and PA 32:0 (down) measured by shotgun lipidomics from human monocyte-derived macrophages with control or CRISPR-mediated SCD1 deletion, stimulated with or without human IL-4 for 48 h. H. Flow cytometry plots of quiescent BMDMs (NT) and BMDMs stimulated with IL-4 ± PA32:0 (10 μM) as indicated for 48 h. Bar graph (right down) depicts percent of double positive (CDCD301+/RELMα+) BMDMs. I. Flow cytometry plots of quiescent BMDMs (NT) and BMDMs stimulated with IL-4 ± PA32:0 (10 μM) as indicated for 48 h. Bar graph (right down) depicts percent of double positive (CDCD301+/Arg1+) BMDMs. J. qPCR analysis of select AAM-associated genes of quiescent BMDM (NT) or BMDMs stimulated with IL-4 ± PA 32:0 (10 μM) for 48 h. K. qPCR analysis of indicated pro-inflammatory genes of quiescent BMDM (NT) or BMDMs stimulated with IL-4 ± PA 32:0 (10 μM) for 48 h. Lipidomics experiments are from four biologic replicates per experimental condition. Heatmap scales are z-score for number of deviations away from the row mean. Rows are clustered using correlation distance and complete linkage. Gene expression and flow cytometry studies are from three biologic replicates per experimental condition and are representative of greater than three experiments. Bar graphs data are presented as mean ± SEM. *p < 0.05; **p < 0.01, ***p < 0.001.

We performed similar lipidomic studies on WT macrophages polarized with IL-4 alone or with IL-4/SCDi. These lipidomics studies largely phenocopied the results of SCD KO macrophages, with a marked accumulation of PAs, DAGs, LPCs, and HexCers when SCD was inhibited, which was normalized to control levels with oleate supplementation (Fig. 4d-f, S4b). Lipidomic time course studies showed that inhibition of SCD did not alter the pool of saturated PAs (PA32:0, PA34:0) at 6 or 12 h timepoints. However, we found substantial PA accumulation at 24 h which continued to increase achieving a nearly 100-fold increase at the 48 h time point (Fig. S4c,d). Importantly, CRISPR-deletion of *SCD1* in human macrophages resulted in the accumulation of saturated PAs in response to IL-4 treatment (Fig. 4g), indicating that inhibiting *de novo* synthesis of MUFAs in human macrophages has a similar impact on the lipidomic reprogramming during AAM polarization.

Several enzymatic pathways can contribute to the homeostasis of PAs in cells. These include de novo synthesis of PAs, the phosphorylation of diacylglycerol (DAG) to form PA by diacylglycerol kinase (DGK) and the remodeling of other phospholipids (e.g., PC)[26]. To gain a better understanding of how PA accumulation was occurring when SCD activity was lost, we looked at PA levels in IL-4 activated macrophages that had ACC1 or FASN inhibited. Analysis of lipidomic data on AAM macrophages treated with either ACCi or FASNi revealed no difference in PA levels when compared to WT AAMs (Fig. S4e, f). Thus, we conclude that the accumulation of saturated PA species (e.g., 32:0, 34:0) results from the specific disruption of MUFA synthesis.

Next, we tested if treating WT (SCD-sufficient) macrophages with PA would attenuate AAM polarization. To that end, we treated cultures of IL-4-stimulated WT macrophages with exogenous PA 16:0_16:0 (PA 32:0; 10μM). Lipidomics showed that cellular PA levels in PA32:0-treated macrophages reached levels similar to SCD-inhibited macrophages (Fig. S4g), without altering cellular DAGs, LPCs, or HexCers (Fig. S4h). Remarkably, we found that PA treatment during AAM polarization markedly decreased the frequency of CD301+/RELMa+ or CD301/Arg1+ macrophages (Fig. 4h, i). Accordingly, we found that PA32:0 treatment blunted the upregulation of AAM-associated genes (*Arg1, Ym1, Mgl2,* and *Retnla*) (Fig. 4j). Notably, we did not observe an increase in pro-inflammatory gene expression when PA was added to IL-4-treated macrophages (Fig. 4k), suggesting that PA’s ability to limit AAM polarization occurs independent of the pro-inflammatory component of SCD inhibition. Accordingly, we asked if PAs would limit pro-inflammatory responses. To address this, we stimulated WT macrophages with TLR2 agonist PAM3CysK alone (50 ng/mL) or with exogenous PA 16:0_16:0 (PA 32:0; 10μM). Importantly, we found that supplementing PA to TLR2-activated macrophages maintained or increased expression of pro-inflammatory genes tested (Fig. S4i). Thus, we conclude that PA accumulation restrains AAM polarization without inducing inflammation and does not limit polarization into pro-inflammatory states. Therefore, PA accumulation acts as a lipid signal that restricts macrophage plasticity.

### PA accumulation attenuates AAM polarization by altering the mTOR signaling axis

PAs have well-defined roles as second messengers[27]. One target of PA is mTOR complex 1 (mTORC1) via direct binding to the FKB domain[28]. Heightened mTORC1 signaling has been shown to negatively regulate AAM polarization[29, 30], leading us to hypothesize that SCD inhibition and the resultant accumulation of PA would activate mTORC1 signaling. To test this hypothesis initially, BMDMs were treated with IL-4 alone, PA32:0 (10 μM) alone, or in combination. 48 h after treatment, whole-cell lysates were collected and immunoblots were performed to assess the phosphorylation states of the mTORC1 target S6 kinase (Thr 389) and the mTORC2 target AKT (Ser 473). We observed that IL-4 alone increased p-AKT (Ser 473), but p-S6 kinase levels were similar to control, consistent with literature describing heightened mTORC2 signaling in AAMs[31] (Fig 5a). Consistent with a role for PA regulating mTOR, PA32:0 treatment of IL-4-activated macrophages resulted in heightened phospho-S6 kinase and reduced p-AKT (Fig. 5a). Thus, accumulation of saturated PAs alters mTOR signaling during AAM polarization.

**Figure 5.**
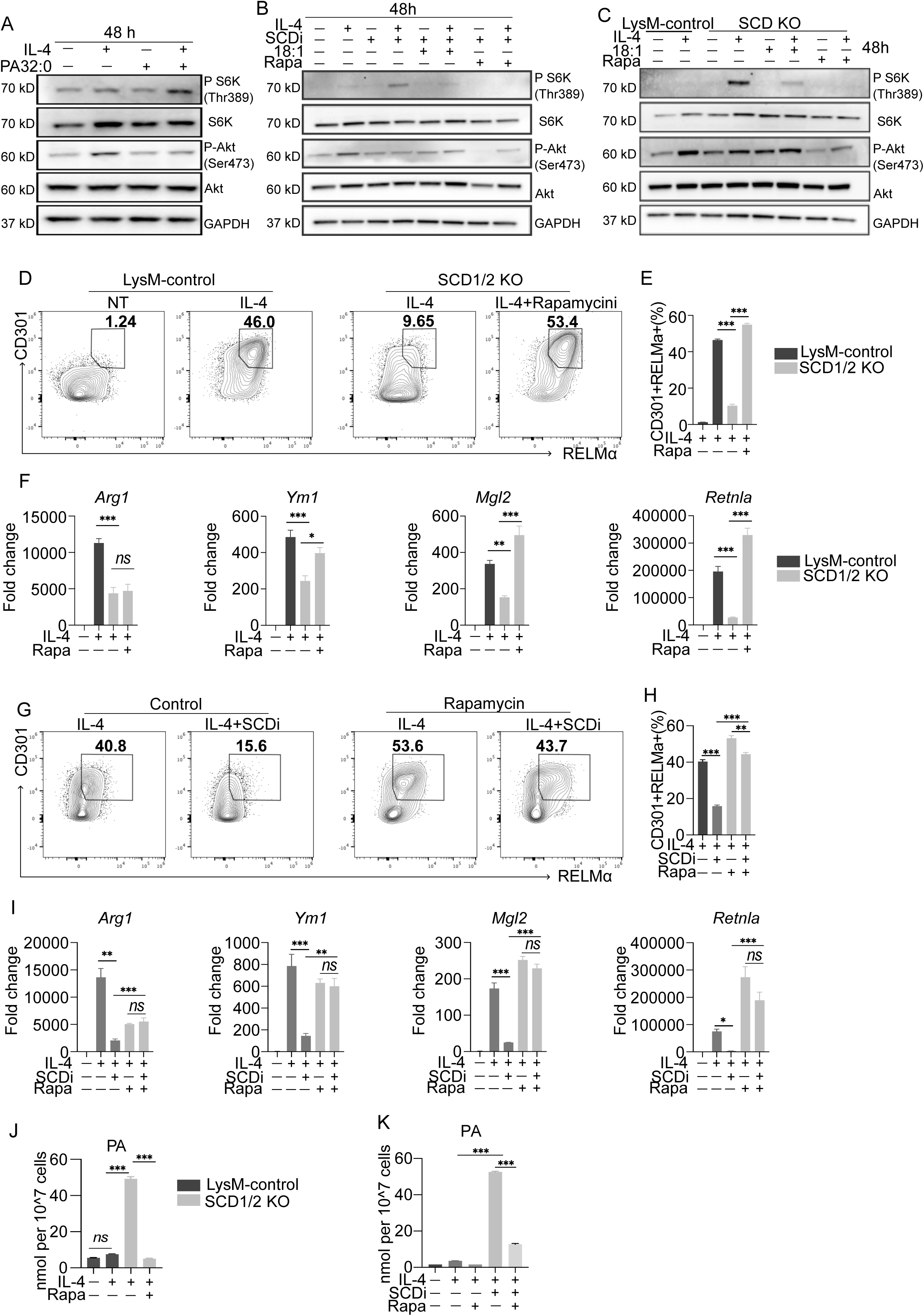
PA-mTORC1 signal initiated by loss of MUFA synthesis restricts AAM polarization. A. Western blot analysis of total and phosphorylated S6 kinase and Akt from quiescent BMDMs (NT) or BMDMs stimulated with IL-4 ± PA32:0 (10 μM) or PA 32:0 (10 μM) alone as indicated for 48 h. B. Western blot analysis of total and phosphorylated S6 kinase and Akt from quiescent BMDMs (NT) or BMDMs stimulated with IL-4 or IL-4+SCDi ± 18:1(25 μM) or IL-4+SCDi ± Rapamycin (10 nM) or SCDi ± 18:1(25 μM), or SCDi ± Rapamycin (10 nM) as indicated for 48 h. C. Western blot analysis of total and phosphorylated S6 kinase and Akt from LysMCre control and LysMCre-SCD1/2^fl/fl^ BMDMs stimulated with or without IL-4, or LysMCre-SCD1/2^fl/fl^ BMDMs stimulated IL-4 ± 18:1 (25 μM), or IL-4 ± Rapamycin (10 nM) as indicated for 48 h. D. Flow cytometry plots from LysMCre control BMDMs stimulated with or without IL-4 and LysMCre-SCD1/2^fl/fl^ BMDMs stimulated with IL-4 ± Rapamycin (10 nM) as indicated for 48 h. E. Bar graph depicts percent of double positive (CDCD301+/RELMα+) BMDMs as above. F. qPCR analysis of select AAM-associated genes from LysMCre control BMDMs stimulated with or without IL-4 and LysMCre-SCD1/2^fl/fl^ BMDMs stimulated with IL-4 ± Rapamycin (10 nM) as indicated for 48h. G. Flow cytometry plots from BMDMs stimulated IL-4 ± SCDi or IL-4 ± SCDi with Rapamycin (10 nM) as indicated for 48 h. H. Bar graph depicts percent of double positive (CDCD301+/RELMα+) BMDMs as above. I. qPCR analysis of select AAM-associated genes from quiescent BMDMs (NT) or BMDMs stimulated with IL-4 ± SCDi or IL-4 ± SCDi with Rapamycin (10 nM) as indicated for 48h. J. Total phosphatidic acid (PA) measured by shotgun lipidomics from LysMCre control BMDMs stimulated with or without IL-4 and LysMCre-SCD1/2^fl/fl^ BMDMs stimulated with IL-4 ± Rapamycin (10 nM) as indicated for 48 h. K. Total phosphatidic acid (PA) measured by shotgun lipidomics from quiescent BMDMs (NT) or BMDMs stimulated with IL-4 ± SCDi or IL-4 ± SCDi with Rapamycin (10 nM) as indicated for 48 h. Lipidomics experiments are from four biologic replicates per experimental condition. Heatmap scales are z-score for number of deviations away from the row mean. Rows are clustered using correlation distance and complete linkage. Gene expression and flow cytometry studies are from three biologic replicates per experimental condition and are representative of greater than three experiments. Bar graphs are presented as mean ± SEM. *p < 0.05; **p < 0.01, ***p < 0.001.

Next, we asked if loss of SCD alters mTOR signaling. To test this, WT macrophages were stimulated with IL-4 alone or in combination with SCDi, SCDi+18:1, or SCDi+rapamycin (an mTORC1 inhibitor). Similar to results with PA supplementation, treatment with SCDi in combination with IL-4 resulted in heightened phospho-S6 kinase and reduced phospho-AKT. Importantly, the addition of oleate, markedly reduces phospho-S6 kinase to levels near that of rapamycin treatment (Fig. 5b). Despite this reduction in mTORC1 signaling by oleate or rapamycin, phospho-AKT remained well below that of IL-4 treatment alone (Fig. 5b). Immunoblots on lysates from IL-4-activated *LysM*-control and SCD1/2 KO macrophages showed a similar pattern where phospho-S6 kinase levels were elevated and phospho-AKT levels reduced (Fig. 5c). Again, the addition of oleate to cultures markedly reduces phospho-S6 kinase and partially restored phospho-AKT (Fig. 5c). These data show that genetic or pharmacologic disruption of MUFA biosynthesis perturbs the mTOR signaling axis during AAM polarization.

We reasoned that inhibiting mTOR signaling would restore AAM polarization in SCD deficient macrophages. To test this idea, *LysM*-control and SCD1/2 KO BMDMs were activated with IL-4 alone or in combination with rapamycin. As expected, the frequency of CD301+/RELMα+ or CD301/Arg1+ macrophages was markedly decreased in SCD1/2 KO macrophages (Fig. 5d,e and S5a,b). Remarkably, the addition of rapamycin to SCD1/2 KO cultures was able to normalize the upregulation of CD301+/RELMα+ or CD301/Arg1+ macrophages (Fig. 5d,e and S5a,b). Correspondingly, rapamycin treatment largely restored upregulation of AAM-associated genes *Ym1, Mgl2,* and *Retnla*, although *Arg1* was not rescued (Fig. 5f). We also found that rapamycin restored phenotypic differentiation (e.g., CD301+/RELMα+) and the upregulation of AAM gene expression (*Ym1, Mgl2,* and *Retnla)* in WT macrophages treated with SCDi (Fig. 5g-i and S5 c,d). Notably, lipidomics revealed that rapamycin blunted the aberrant accumulation of PA in SCD loss-of-function macrophages (Fig. 5j,k and S5 e,f). Together, these data support a model where mTORC1 inhibition restores AAM polarization in SCD loss-of-function macrophages by blunting PA accumulation.

### Disrupting the SCD–PA lipid regulatory axis attenuates pathogen-directed alternative activation of macrophages

Our data support a model where loss of SCD and the resulting accumulation of saturated phosphatidic acids attenuate the acquisition of an alternatively activated cell state. The type III strain CEP of the intracellular parasite *T. gondii* converts infected macrophages into an alternatively activated phenotype by the parasite effector protein ROP16[32]. This conversion to AAMs reduces pro-inflammatory anti-microbial responses and facilitates the lifecycle of this pathogen. Mechanistically, ROP16 activates STAT6 to drive macrophage conversion to AAMs, independent of IL-4 or IL-13 signaling[32]. Thus, we asked if targeting the SCD-PA regulatory node might interfere with the pathogenic conversion of AAMs by *Toxoplasma* during infection.

Dose response curves (0.5-1.5 MOI) with *Toxoplasma* confirmed that infection of BMDMs robustly induces the AAM gene expression program at 24 and 48 h post parasite challenge (Fig. S6a). Subsequently, we performed all *in vitro* infection experiments using an MOI of 1.5. Notably, gene expression studies showed that *Toxoplasma* infection upregulates *Scd1* and *Scd2* (Fig. 6a). In contrast, *Fasn* levels did not change with parasite infection. We then infected *LysM*-control and SCD KO macrophages to assess AAM conversion. As reported, infection of control macrophages resulted in robust upregulation of AAM-associated genes *Arg1, Ym1, Mgl2,* and *Retnla* (Fig. 6b)[32]. In contrast, loss of SCD attenuated upregulation of AAM genes. Correspondingly, we observed a decrease in parasite burden in SCD KO macrophages as measured by the *Toxoplasma 529bp* repeat element (Fig. 6c). We also performed studies with SCD inhibitors on WT macrophages challenged with *Toxoplasma*. We found a near-identical ability to block the upregulation of the AAM program during infection and a reduced pathogen load (Fig. S6b,c).

**Figure 6.**
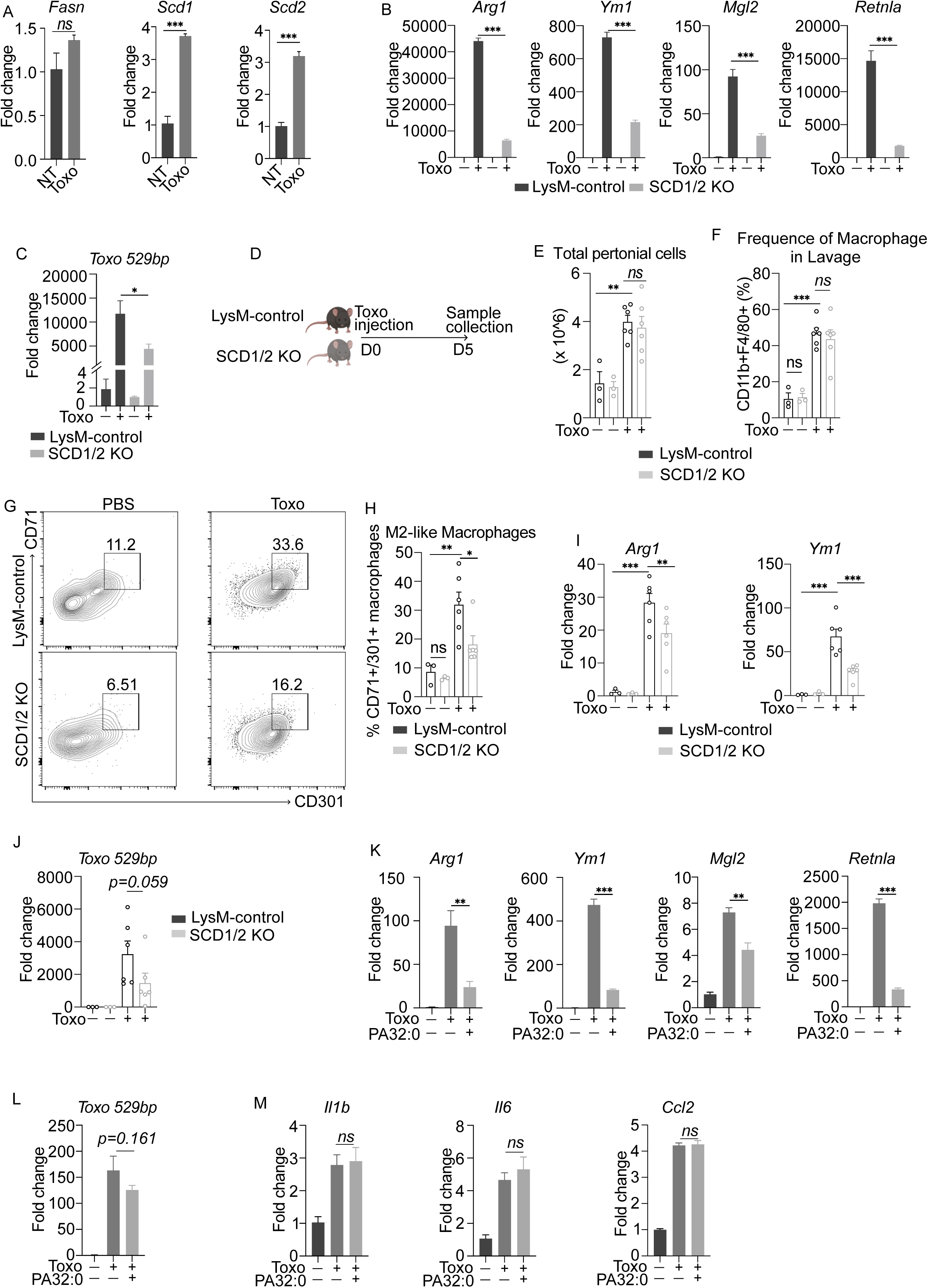
SCD-PA lipid regulatory axis is essential for the conversion of macrophages into AAM during Toxoplasma infection in vitro and in vivo. A. qPCR analysis of *Fasn*, *Scd1*, *Scd2* from quiescent BMDMs (no infection) or BMDMs infected with *Toxoplasma* (MOI=1.5) as indicated for 48h. B. qPCR analysis of select AAM-associated genes from LysMCre-control (WT) and LysMCre-Scd1/2^fl/fl^ BMDMs infected with *Toxoplasma* (MOI=1.5) as indicated for 48h. C. qPCR analysis of *Toxoplasma 529-bp* repeat element (*Toxo 529bp*) from LysMCre-control (WT) and LysMCre-Scd1/2^fl/fl^ BMDMs infected CEP Toxoplasma (MOI=1.5) as indicated for 48h. D. Simplified schematic of the *Toxoplasma in vivo* infection study. E. Bar graph depicts the total number of peritoneal cells collected by PBS lavage from LysMCre-control and LysMCre-Scd1/2^fl/fl^ mice infected with 1000 parasites for 5 days. F. Bar graph depicts percent of double positive (CD11b+/F4/80+) gating from CD45+ peritoneal cells from LysMCre-control and LysMCre-Scd1/2^fl/fl^ mice infected with 1000 parasites for 5 days. G. Flow cytometry plots of peritoneal macrophages collected from LysMCre-control and LysMCre-Scd1/2^fl/fl^ mice infected with 1000 parasites for 5 days. H. Quantification of total double positive (CD301+/CD71+) peritoneal macrophages in the mice treated above. I. qPCR analysis of select AAM-associated genes from peritoneal cells collected by PBS lavage from LysMCre-control and LysMCre-Scd1/2^fl/fl^ mice infected with 1000 parasites for 5 days. J. qPCR analysis of Toxoplasma gondii *529-bp repeat element* (*Toxo 529bp*) in peritoneal cells collected by PBS lavage from LysMCre-control and LysMCre- Scd1/2^fl/fl^ mice infected with 1000 parasites for 5 days. K. qPCR analysis of select AAM-associated genes from quiescent BMDMs (no infection) or BMDMs infected with *Toxoplasma* (MOI=1.5) ± PA 32:0 (10 μM) for 48 h. L. qPCR analysis of the *Toxoplasma 529-bp* repeat element (*Toxo 529bp*) from quiescent BMDMs (no infection) or infected BMDMs (MOI=1.5) ± PA 32:0 (10 μM) for 48 h. M. qPCR analysis of indicated pro-inflammatory genes from quiescent BMDMs (no infection) or infected BMDMs (MOI=1.5) ± PA 32:0 (10 μM) for 48 h. Gene expression and flow cytometry studies are from three biologic replicates per experimental condition and are representative of greater than three experiments. In vivo experiments are representative of n = 6 mice per group repeated at least two times. Bar graphs are presented as mean ± SEM. *p < 0.05; **p < 0.01, ***p < 0.001.

Next, we challenged *LysM-*control and SCD1/2 KO mice i.p. with *Toxoplasma* (1000 parasites/mouse; Fig. 6d). No differences in body weight between *LysM*-control and SCD1/2 KO mice was noted on day 5 of *Toxoplasma* infection (Fig. S6d). As expected, infected *LysM*-control mice had significant splenomegaly, whereas infected *LysM*-*Scd1/2^fl/fl^*(SCD1/2 KO) mice did not have splenomegaly (Fig. S6e). Peritoneal lavage was collected for flow cytometry and gene expression studies to assess macrophage phenotype and parasite burden. There were no differences in the total number of cells or peritoneal macrophages (CD11b+/F4-80+) in the lavage fluid of *LysM*-control and SCD1/2 KO mice (Fig. 6e,f). However, FACS analysis revealed a decrease in the frequency of AAMs (identified by CD71+/CD301+) from the lavage fluid of infected SCD1/2 KO mice relative to *LysM-*control mice (Fig. 6g,h). Gene expression studies of peritoneal macrophages from lavage fluid also showed upregulation of *Arg1* and *Ym1* in *LysM-*control mice, which was attenuated with loss of SCD (Fig. 6i). *Mgl2* and *Retnla* were downregulated irrespective of genotype at this time point (Fig. S6f), and correspondingly, no difference in intracellular RELMα protein was seen (Fig. S6g). Although the difference in parasite burden between genotypes was not statistically significant, as measured by the *Toxo 529bp*, there was a clear trend toward decreased parasite burden (Fig. 6j).

Having established that loss of SCD attenuates Toxoplasma-mediated conversion of AAMs in vitro and in vivo, we next determined how SCD inhibition altered the lipidome of infected macrophages. To that end, BMDMs were infected with *Toxoplasma* (MOI 1.5) +/- SCDi (10nM). After 48 h, macrophages were collected, lipids extracted and lipidomics performed. Analysis of lipidomics data revealed that *Toxoplasma* markedly reshaped the macrophage lipidome (Fig. S6h), much of which was dependent on SCD activity. Inspection of lipids accumulated in SCD-inhibited macrophages revealed that PA species were the most changed (Fig. S6i; brown box; see Supp Table 2 for full lipidomic data), paralleling our observations in IL-4 activated macrophages. These data led us to ask if treating macrophages with PA alone would be sufficient to attenuate AAM conversion by *Toxoplasma* infection. To test this, WT macrophage cultures were concurrently infected (MOI 1.5) and treated with 32:0 PA (10 μM) for 48 h. Paralleling our results in SCD KO or inhibition, we found that PA treatment markedly attenuated parasite-driven upregulation of AAM genes (Fig.6k). Of importance, this block in upregulation of the AAM program did not influence parasite burden at this time point (Fig. 6l), nor did it augment pro-inflammatory genes (Fig. 6m, S6j). Thus, accumulation of saturated PAs in macrophages, can disrupt *Toxoplasma*-driven AAM conversion independent of PAs role in fostering inflammation in TLR-activated macrophages. Together, these data provide proof-of-concept evidence that perturbing the SCD-PA signaling axis in macrophages disrupts pathogen-driven lipid metabolic reprogramming and the accompanying AAM conversion.

## DISCUSSION

In this study, we identify stearoyl-CoA desaturase (SCD) enzymes as key regulators of macrophage cell state and functional plasticity. Our goal was to determine how and why the macrophage lipidome is reprogrammed in response to signals that instruct alternative activation. We learned that a considerable portion of the lipidome is reshaped during AAM polarization to IL-4, and that much of this reprogramming is dependent on *de novo* lipogenesis (DNL). We found that lipids accumulated during AAM polarization were enriched for monounsaturated fatty acid chains, suggesting that a major purpose of DNL flux is to provide substrate for the SCD enzymes. Consistent with this, loss of SCD and MUFA synthesis impaired the AAM cell state at the epigenetic, transcriptional, and phenotypic levels. Interestingly, inhibition of DNL enzymes ACC1 and FASN did not block AAM polarization under these conditions. Therefore, an imbalance between the synthesis of saturated and monounsaturated fatty acids (e.g., the desaturation index), rather than loss of DNL itself, is responsible for disrupting AAM polarization. Furthermore, pharmacologic inhibition or genetic deletion of SCD rendered macrophages resistant to *Toxoplasma*-induced conversion to an alternatively activated state. Thus, we conclude that *de novo* MUFA synthesis serves as a lipid metabolic checkpoint that ensures macrophages remain responsive to anti-inflammatory or pro-resolving information.

The mechanisms by which SCD enzymes regulate macrophage state remain incompletely understood. We propose that at least two distinct molecular circuits are activated when cellular MUFA synthesis is restricted. In this study, we show that disruption of SCD leads to the aberrant accumulation of saturated phosphatidic acids (PAs), which interferes with stable acquisition of the AAM program. Consistent with this, provisioning saturated PAs to cultures disrupted AAM polarization of wild-type macrophages, phenocopying SCD loss. Mechanistically, we found that accumulated saturated PAs activate mTORC1 signaling, presumably through its ability to interact the FKB domain[33, 34]. Consequently, the balance between mTORC1 and mTORC2 activity is altered when SCD is inhibited or PA accumulates during AAM polarization. The mTOR signaling pathway is a well-established regulator of macrophage fate, where persistent activation of mTORC1, or mTORC2 loss, has been shown to attenuate AAM polarization[29, 30]. Consistent with a role for PA and mTOR signaling in restricting AAM polarization when MUFA synthesis is lost, pharmacologic inhibition of mTORC1, using rapamycin, largely restored AAM polarization potential in SCD-deficient macrophages. Of importance, accumulation of saturated PAs in TLR-activated macrophages did not impair pro-inflammatory responses or attenuate M1 polarization. Thus, the PA–mTORC1 axis appears to selectively restrict acquisition of an AAM cell state rather than globally disrupting macrophage polarization or identity.

The observation that addition saturated PAs to IL-4-activated macrophages did not induce an inflammatory response suggests that a second lipid signaling axis is responsible for heightened pro-inflammatory cell state of SCD-deficient macrophages. We recently showed that the loss of SCD2 in the context of TLR2 activation leads to the accumulation of saturated very long-chain (VLC) ceramides (24:0 Cer). The accumulation of VLC ceramides was dependent on *de novo* synthesis by ceramide synthase enzymes and resulted in increased nuclear NF-κB, which consequently prolonged inflammation[16]. Consistent with this mechanism, lipidomic analysis of IL-4–activated SCD-deficient macrophages revealed a modest increase in VLC ceramides. We also found increases in DAGs and LPCs when SCD was inhibited during AAM polarization. LPCs can promote inflammation by serving as intermediates for the generation of platelet-activating factor (PAF) and lysophosphatidic acid (LPA)[35, 36]. Likewise, DAGs act as a signaling molecule that activates protein kinase C, driving inflammatory responses[37]. Future studies will be needed to determine which of these lipids signaling pathways contributes to redirecting SCD-deficient macrophages into an altered cell state during AAM polarization.

Finally, it remains unclear why the accumulation of saturated PAs selectively restricts AAM polarization while preserving macrophage responsiveness to pro-inflammatory cues. One plausible explanation is that rapid elevations in PA serve as an intrinsic danger signal that shapes host defense against intracellular pathogens. Consistent with this idea, viral and intracellular parasitic infections have been shown to increase PA levels to facilitate their lifecycle[38–40]. Likewise, plants rapidly synthesize PAs in response to microbial invasion or damage, triggering innate immune components such as the oxidative burst[41–43]. The intracellular parasite *T. gondii* is remarkably efficient at infecting all mammals, resulting in lifelong infections of hosts in upwards of a third of the world’s human population[44–46]. Toxoplasma and other pathogens have been shown to drive the conversion of infected macrophages into an AAM-like state to facilitate pathogen persistence[32, 47]. We found that macrophage infection with the *Toxoplasma* CEP strain markedly upregulated host *Scd1* and *Scd2* expression, suggesting that heightened MUFA synthesis, and consequently maintaining low intracellular PA levels, supports parasite lifecycle. Consistent with this concept, increasing PA levels, either by targeting SCD or treating with 32:0 PA, attenuated macrophage conversion to the AAM state. We think it likely that other lipids perturbed by SCD inhibition during *Toxoplasma* infection will also foster pro-inflammatory responses to further limit pathogen lifecycle. It will be of interest in future studies to determine whether restricting AAM polarization as a consequence of SCD inhibition or PA accumulation will translate into enhanced host immunity or perturbed *Toxoplasma* persistence.

In conclusion, our findings advance our understanding of how lipid metabolic reprogramming governs the acquisition of macrophage effector states and establish SCD enzymes as pivotal regulators of macrophage plasticity.

## ADDITIONAL INFORMATION

### Data availability

RNA-Seq and ATAC-Seq data is deposited at Gene Expression Omnibus (GEO) and will be publicly available as of the date of publication.

Lipidomics data is deposited at EMBL’s European Bioinformatics Institute (EMBL_EBI) Metabolights repository and will be publicly available as of the date of publication.

## Acknowledgments

This work was supported by funding K08 HL179476 (RLW), R01 HL175240 (PT), R01 DK133344 (CFS), R01 NIH AI123360 (PJB), Ruth L. Kirschstein National Research Service Award AI007323 (ACT), Hughes Medical Institute Hanna H. Gray Fellowship. (AGY), R01 HL157710 (SJB). We thank Yu Qiao, Kelly Kennewick, and Alexandra Kuhlmann for their thoughtful review and input on the manuscript.

## Author Contributions

T.H., and R.L.W. conceptualized, designed and/or implemented experiments, analyzed data, and manuscript construction. AGY, AB, JQ, ACT, JNU, and W-Y H conceptualized, designed and/or implemented experiments, and/or analyzed data. C.F.S., P.J.B., and P.T. provided resources, supervision, and/or contributed to conceptualization. S.J.B. provided resources, supervision, and contributed to conceptualization/design of experiments, and manuscript construction.

## Declaration of interests

The authors declare no competing interests.

## METHODS AND MATERIALS

REAGENT or RESOURCE

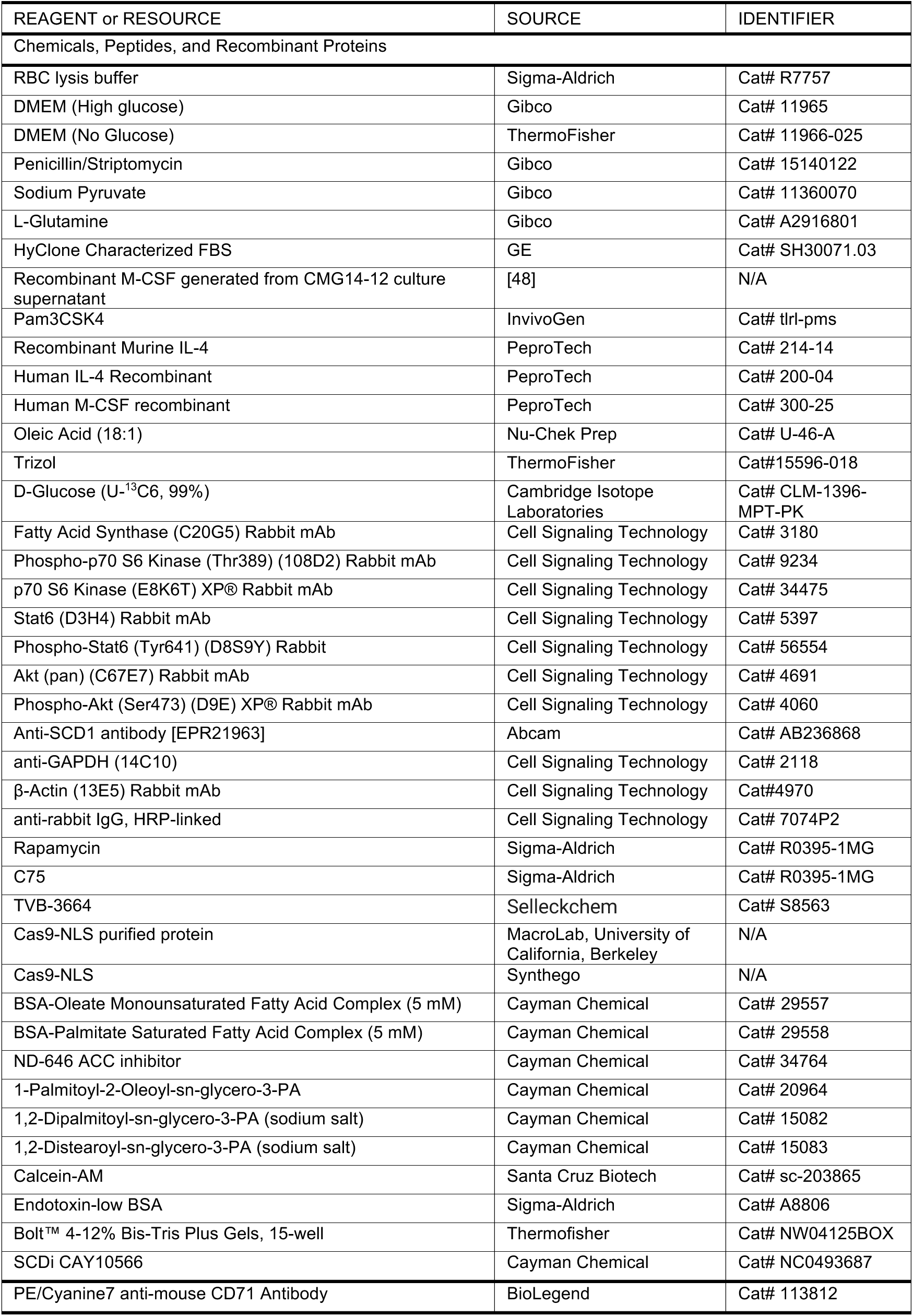
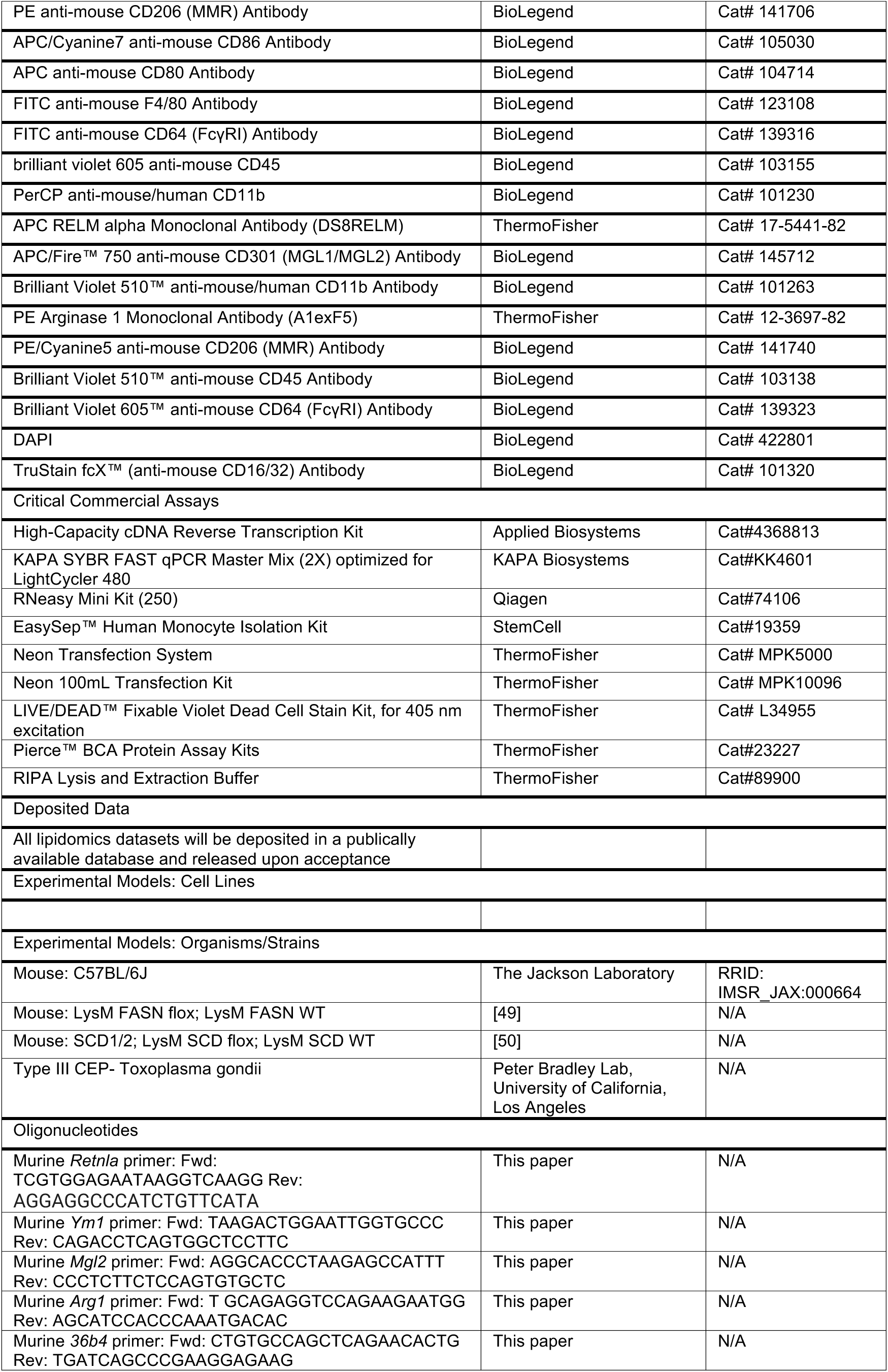
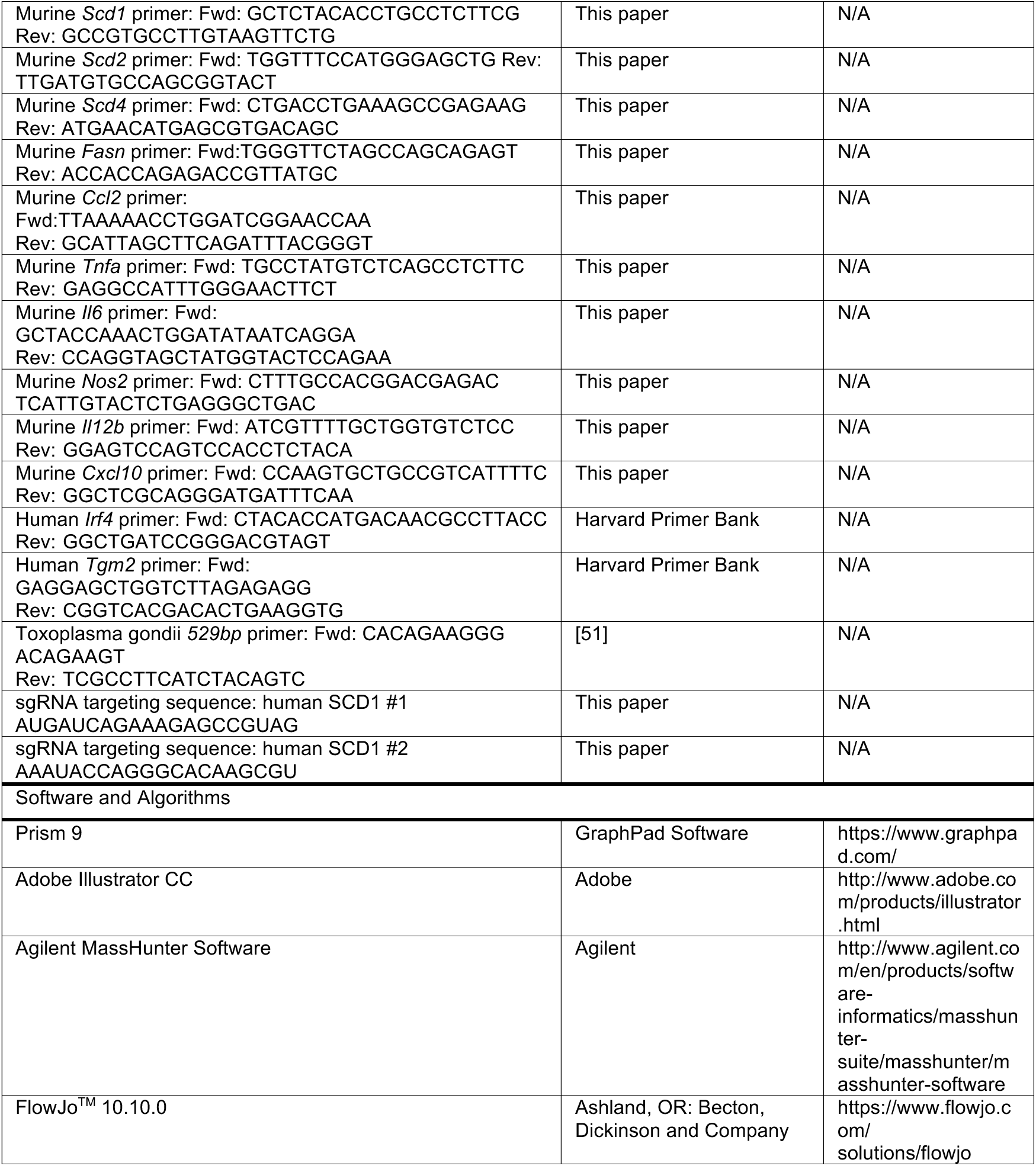

## METHODS

### Isolation of bone marrow-derived macrophages (BMDMs)

For all experiments involving BMDMs except for flux analysis, cells were cultured in either DMEM supplemented with 5% or 10% heat-inactivated fetal bovine serum (FBS)(designated as 5% or 10% BMDM media) plus 2 mM L-glutamine, 100 units/mL, 100 μg/mL penicillin/streptomycin, 500 µM sodium pyruvate, and 5% v/v conditioned media containing macrophage colony stimulating factor (M-CSF) produced by CMG cells to induce differentiation to BMDMs.

Bone marrow cells were isolated from femurs of male C57BL/6 mice or FASN KO and SCD KO mice. Cells were treated with RBC lysis buffer to remove red blood cells for three minutes, centrifuged at 365g for 5 minutes, and resuspended in 10% BMDM medium. Cells were maintained at 37°C in a humidified 5% CO_2_ incubator. BMDMs were differentiated for 6 days prior to experiments, and medium was changed at Day 4 of differentiation.

### Lipidomics analysis of BMDM

Day 6 BMDMs were seeded at 9×10^5^ cells/well in 6-well plates in 10% BMDM media. 48 h later, BMDM were mock-treated or co-treated with ligands/inhibitors/lipids as indicated in 5% BMDM media for 48 h. Cell number was assessed as described below prior to sample collection. All lipidomics analysis were conducted by UCLA lipidome core facility center.

### Isotope labeling analysis of BMDM

Day 6 BMDMs were seeded at 1×10^5^ cells/well in 24-well black plates in 10% BMDM media. 48 h later, BMDM were mock-treated or co-treated with inhibitors for SCD,FASN,ACC plus 20 ng/mL IL-4 in DMEM (no glucose) supplemented with 10 mM [^13^C_6_]-glucose and 10 mM unlabeled glucose (i.e., 50/50 mixture of heavy and light glucose), 5% FBS, 2 mM L-glutamine, 100 units/mL penicillin, 100 μg/mL streptomycin, and 5% v/v conditioned media for 48 h. Isotopomer spectral analysis (ISA) was conducted using an Agilent 5975C mass spectrometer coupled to a 7890 Gas Chromatograph as previously described[52]. Data were normalized by cell count prior to sample collection. For collection with guanidine HCl, cells were treated with 50 uL of 6 M aqueous guanidine HCl and transferred to glass tubes for derivatization with an additional 100 uL of 3 M methanolic guanidine HCl. Samples were prepared alongside internal standard curve samples made up of FAMEs (Nu-Chek Prep).

### Cell counts and normalization

For both flux and lipidomics experiments, prior to sample collection, 1.25 mM Calcein-AM dissolved in glucose-free DMEM (final concentration; Santa Cruz Biotechnologies) was added to each well and the plates were then imaged on a Molecular Devices ImageXpress XL (CNSI, UCLA). 20 high magnification fluorescence images were captured for each well (21.83% of total well surface area) using a 10x objective (Nikon Plan Fluor, 0.3 NA). Cell number was assessed using MetaXpress Software with Powercore using the multi-wavelength cell scoring module.

### Gene expression analysis

For all experiments involving gene expression analysis, day 6 BMDMs were seeded at 2.5×10^5^ cells/well in 12-well plates in 10% BMDM media. 48 h later, BMDM were mock-treated or co-treated IL-4 (20ng/mL) with inhibitors/lipids in 5% BMDM media as indicated in the figure legend. Cells were collected in TRIzol and RNA was extracted according to manufacturer’s protocol. cDNA was synthesized using 700 ng RNA per reaction with high-capacity cDNA reverse transcription kit. KAPA SYBR FAST qPCR Master Mix (2X) kit and a LightCycler 480 were used for quantitative RT-PCR. Fold change related to the control group was calculated using 2^ΔΔCP^ method with *36b4* as the reference gene.

### Western blot analysis

Samples were lysed in RIPA Buffer (ThermoFisher) and normalized by BCA Protein Assay Kits (ThermoFisher) before 1:1 dilution with 2x Laemmli loading buffer. Protein extracts were separated on gradient 4% to 12% Bis–Tris SDS-PAGE gel (Invitrogen) and then transferred to a nitrocellulose membrane (Amersham) or PVDF membrane (Millipore Sigma). After blocking for 1 hour in a TBS containing 0.1% Tween 20 (TBST) and 5% nonfat milk, the membrane was probed with indicated antibodies diluted into TBST with 5% milk (1:1000) overnight. Membranes were washed 4x with TBST, followed by a 60-minute room temperature incubation with secondary antibodies (1:2000) conjugated to horseradish peroxidase 3 diluted in TBST plus 5% milk. Membranes were washed as before and then developed using Pierce ECL2 detection kit and imaged with Typhoon.

### Flow cytometric analysis

For all BMDM polarization experiments, day 6 BMDMs were seeded at 3×10^5^ cells/well in 12-well plates in 10% BMDM media. 48 h later, BMDM were mock-treated or treated with ligands as indicated in individual figures in 5% BMDM media. BMDM were mock-treated or co-treated with inhibitors/lipids as indicated in 5% BMDM media ± 20ng/mL of IL-4. Samples were collected 48 h post simulation. Prior to FACS analysis, BMDMs were detached using 5mM EDTA with cold PBS for 10 mins, followed by two washes using the FACS buffer (PBS supplemented with 2% FBS and 1 mM EDTA). Cells were stained with a LIVE/DEAD fixable violet viability dye (1:1000, 405nm excitation; ThermoFisher) and intubated with TruStain FcX receptor blocker (1:100, BioLegend). Cells were then stained with the indicated fluorochrome-conjugated antibodies (manufacturer, clone) at a 1:200 dilution. All flow cytometry data were captured using Attune NxT Flow Cytometer (ThermoFisher). Compensation and analysis were performed using FlowJo software (BD Biosciences).

### *In vitro Toxoplasma gondii* Infection

Day 6 BMDMs were seeded at 2.5×10^5^ cells/well in 12-well plates in 10% BMDM media. 48 h later, BMDM were either mock-infected or infected with CEP strain *T.gondii* expressing mCherry at a multiplicity of infection (MOI) of 0.5, 1.0, or 1.5, with or without the indicated inhibitor or lipid treatment[53]. Infected cells were cultured in 5% BMDM medium for 24 or 48 hours. Macrophage cultures were harvested in TRIzol for subsequent gene expression analysis.

### *In vivo Toxoplasma gondii* Infection

*LysM* control and LysM Cre-SCD1/2^fl/fl^ mice between the age of 8-10 weeks (sex matched) were intraperitoneally injected (I.P.) with either 200 ul of vehicle (PBS) or 1000 of CEP strain T. gondii expressing mCherry[53]. Mice were euthanized 5 days after infection via isoflurane overdose. Body weight and spleen weight were measured and immune cell population from peritoneal flushes were analyzed by flow cytometry. Briefly, 5 mL of PBS was I.P. injected into each animal using a 27-gauge needle and 3.5 mL of the peritoneal flush was collected using a 23-gauge needle. Cells were first pelleted by centrifugation at 365g for 7 minutes then resuspend in 1 mL of RBC lysis buffer for 3 mins. RBC lysis buffer was subsequently neutralized by adding 1 mL of PBS. Cells were re-pelleted by centrifugation at 365g for 5 minutes and resuspended in 300 ul of FACS buffer before transferring to a 96-well round bottom plate for staining.

### RNA-seq sample preparation and data analysis

Day 6 BMDMs were seeded at 1×10^6^ cells/well in 6-well plates in 10% BMDM media. 48 h later, BMDM were mock-treated or treated with inhibitors or/and IL-4 as indicated in 5% BMDM media. Samples were collected separately at 0, 6-, 12-, 24-, and 48-hours post stimulation. RNA was extracted using QIAzol reagent (Qiagen, USA), and 1 µg of RNA per sample was used to create sequencing libraries with the NEBNext® Ultra™ RNA Library Prep Kit for Illumina® (NEB, USA). Unique index codes were added to distinguish samples. Samples were clustered using the TruSeq PE Cluster Kit v3-cBot-HS (Illumina, USA) and sequenced on a NovaSeq PE150 platform to generate 150 bp paired-end reads. Low-quality reads were removed, and the clean reads were mapped to the mouse genome using HISAT (version 2.0.4). Gene expression was measured as fragments per kilobase of transcript per million mapped reads (FPKM) using HTSeq (version 0.6.1). Differential expression analysis of the four treatments was done with DESeq (version 1.10.1), identifying significantly differentially expressed genes (DEGs) at P<0.05. Pathway analyses were performed using the fgsea package in R and GSEA and MSigDB references pathways.

### ATAC-seq sample preparation and data analysis

ATAC-seq was performed following standard protocols[54]. In summary, 5,000–50,000 cells were washed with cold PBS, lysed in resuspension buffer (RSB) containing detergents (0.1%NP40, 0.1%Tween-20, and 0.01% digitonin), and incubated on ice for 3 minutes. Nuclei were then washed with RSB, collected by centrifugation, and resuspended in a transposition mix containing Tn5 transposase. The samples were incubated at 37°C for 30 minutes with gentle mixing. After DNA purification using a Qiagen MinElute kit, the DNA was amplified with indexed primers, with the optimal PCR cycle number determined by qPCR. Libraries were size-selected with AmpureXP beads and sequenced on an Illumina NextSeq500 platform to produce 75 bp single-end reads. Single-end reads were aligned to the *Mus musculus* mm10 genome using standard methodologies. A sliding window approach using csaw was used to identify windows at least 8-fold above background. Differentially accessible regions (DARs) were then determined using DESeq2 and motif enrichment was subsequently performed using monaLisa. Stability selection (LASSO regularization) was used to detect motif instances in each region and compete motifs features against each other, independent of CpG content and GC enrichment.

### CRISPR-mediated SCD1 deletion in primary human monocyte-derived macrophages

Peripheral blood mononuclear cells (PBMCs) were collected from deidentified healthy donors by the UCLA virology core. CD14+ cell fraction was isolated using magnetic bead negative separation (Human Monocyte EasySep Kit, StemCell). At day zero, monocytes underwent CRISPR deletion as follows. Guides (120pmol), Alt-R electroporation enhancer (9 pmol, IDT), and Cas9- were combined and allowed to form complexes for 10 minutes on ice. Monocytes were then electroporated with CRISPR complexes using a Neon NxT transfection system (ThermoFisher) with a voltage of 1,900V and one pulse of 20ms width. Cells were then allowed to recover for 60 minutes and plated at desired densities (2.5 ×10^5^ cells per 24-well plate, 5 ×10^5^ cells per 12-well plate, 1.5 ×10^6^ cells per 6-well plate). Cells were then differentiated in RPMI with 10% FBS and recombinant human M-CSF (50ug/ml, Peprotech) for 4-days, with the media being changed every other day. On day 4, differentiated hMDMs were stimulated with human IL-4 (50ng/ml,) for 48 h. Samples were collected for subsequent gene expression, protein, and lipidomic analyses.

### Quantification and statistical analysis

All statistical parameters, including the number of replicates (n), can be found in the figure legends. Statistical analyses were performed using Graph Pad Prism 9 software. Data are presented as the mean ± SEM. For all gene expression analysis, two-tailed unpaired Student’s t-test was used. For flux analysis involving only WT BMDMs, statistical significance determined using ordinary one-way ANOVA followed by Dunnett post-hoc multiple comparisons tests. For flux analysis involving, SCD1/2 KO, FASN KO and SCDi/FASNi/ACCi for each fatty acid measured, statistical significance determined using the multiple Student’s t-tests corrected for multiple comparison using Holm-Sidak method, with alpha = 0.05. Computations assume that all rows are sample from populations with the same scatter (SD). Numbers of mice used in cohorts were pre-determined based on prior literature[55] and no exclusion criteria were used.

## SUPPLEMENTAL FIGURE LEGENDS

**Figure S1.**
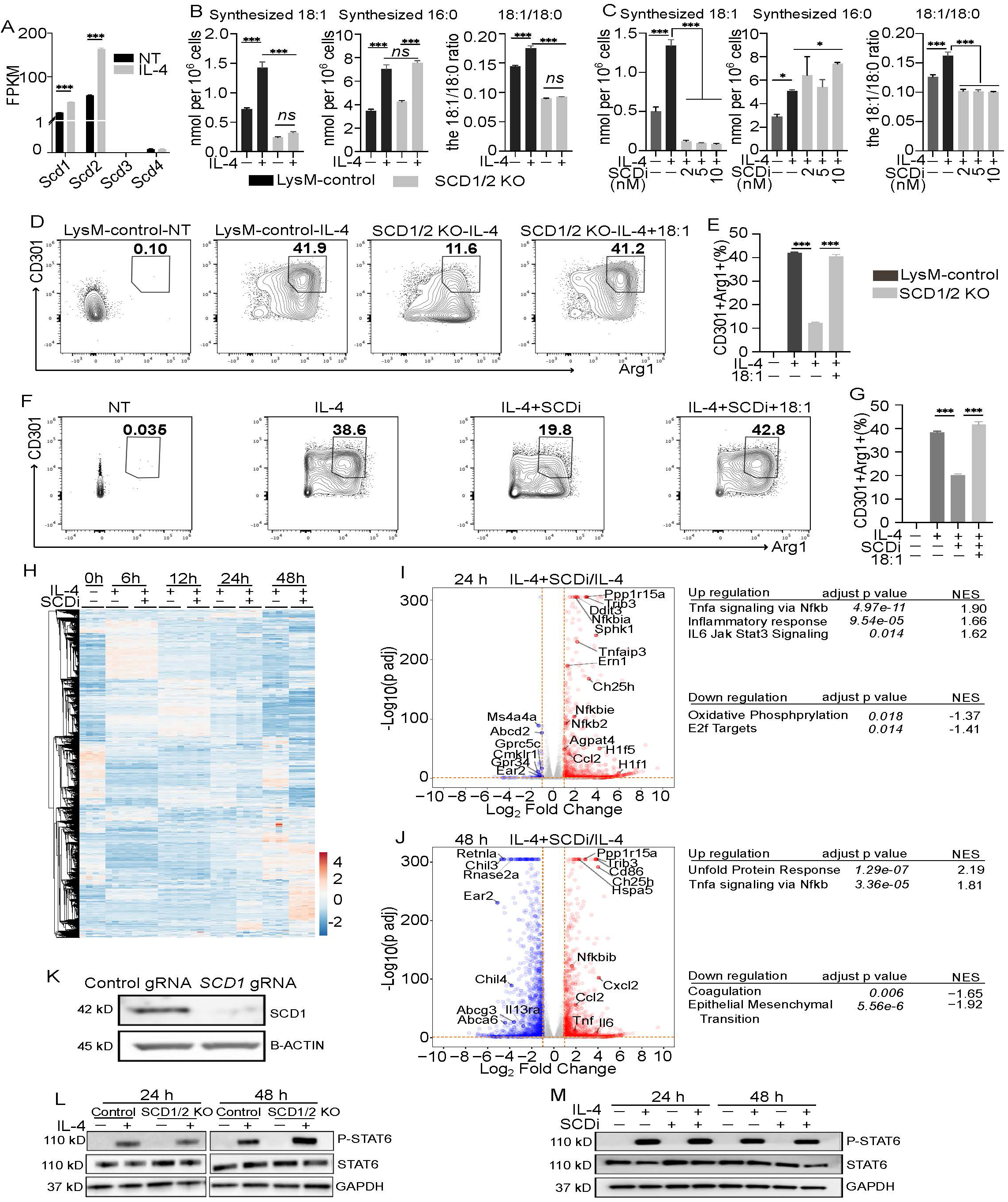
SCD-mediated synthesis of MUFAs is essential for AAM polarization. A. FPKM values of Scd1, Scd2, Scd3, and Scd4 from RNA-Seq studies on quiescent BMDMs (NT) or IL-4 activated BMDMs at 48 h. B. Net synthesized oleic acid (18:1) and palmitic acid (16:0) from LysMCre control and LysMCre-SCD1/2^fl/fl^ BMDMs stimulated with or without IL-4 for 48 h. Fatty acid desaturation index determined by comparing the amount of oleic acid (18:1) to stearic acid (18:0). C. Net synthesized oleic acid (18:1) and palmitic acid (16:0) from quiescent BMDMs (NT) or BMDMs stimulated with IL-4 ± indicated doses of SCDi for 48 h. Fatty acid desaturation index determined by comparing the amount of oleic acid (18:1) to stearic acid (18:0). D. Flow cytometry plots of LysMCre control with or without IL-4 and LysMCre-SCD1/2^fl/fl^ BMDMs stimulated with IL-4 ± 18:1 (25 μM) as indicated for 48 h. E. Bar graph depicts percent of double positive (CDCD301+/Arg1+) BMDMs in D. F. Flow cytometry plots of quiescent BMDMs (NT) and BMDMs stimulated with IL-4 or IL-4+SCDi (Cay10566) ± 18:1(25 μM) as indicated for 48 h. G. Bar graph depicts percent of double positive (CDCD301+/Arg1+) BMDMs in F. H. Heatmap of RNA-seq data of quiescent BMDMs (NT) or BMDMs stimulated with IL-4 ± SCDi for 0h, 6h, 12h, 24 h and 48h. Genes (rows) were clustered according to correlation. I. Volcano plot of differentially expressed genes (DEGs) from RNA-seq analysis of BMDMs stimulated with IL-4 ± SCDi for 24 h. J. Volcano plot of differentially expressed genes (DEGs) from RNA-seq analysis of BMDMs stimulated with IL-4 ± SCDi for 48 h. K. Western blot analysis of SCD1 from control or CRISPR-mediated SCD1 deletion of human monocyte-derived macrophages stimulated with IL-4 for 48 h. L. Western blot analysis of total and phosphorylated STAT6 from LysMCre-control (Control) and LysMCre-SCD1/2^fl/fl^ BMDMs stimulated with or without IL-4 for 24 or 48 h. M. Western blot analysis of total and phosphorylated STAT6 from BMDMs (NT) and BMDMs stimulated with IL-4 ± SCDi or SCDi alone for 24 h or 48 h. All experiments are from four biologic replicates per experimental condition. Heatmap scales are z-score for number of deviations away from the row mean. Rows are clustered using correlation distance and complete linkage. Gene expression and Flow cytometry studies are from three biologic replicates per experimental condition and are representative of greater than three experiments. Bar graphs are presented as mean ± SEM. *p < 0.05; **p < 0.01, ***p < 0.001.

**Figure S2.**
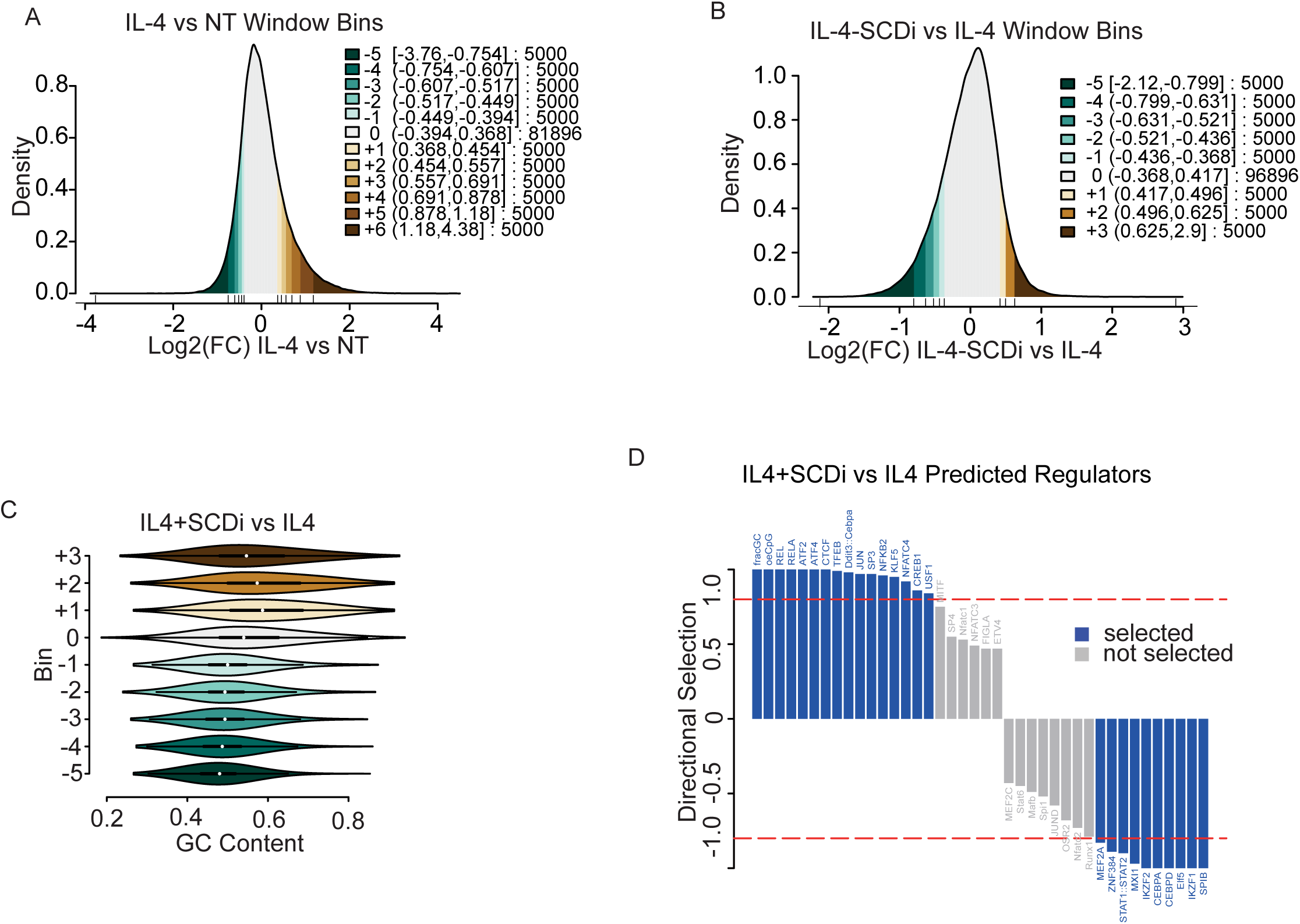
SCD inhibition drives AAM chromatin remodeling independent of potential bias. A. Histogram depicting differentially accessible regions (DARs) for each of the sliding window bins identified using csaw, in quiescent BMDMs (NT) and BMDMs stimulated with IL-4 for 24 h. B. Histogram depicting differentially accessible regions (DARs) for each of the sliding window bins identified using csaw, in BMDMs stimulated with IL-4 ± SCDi for 24 h. C. Plot of GC content in each of the sliding window bins identified in BMDMs stimulated with IL-4 ± SCDi for 24 h. D. De novo motifs identified using LASSO regularization in quiescent BMDMs (NT) and BMDMs stimulated with IL-4 ± SCDi for 24 h. All ATAC-seq and RNA-seq experiments are from three biologic replicates per experimental condition.

**Figure S3.**
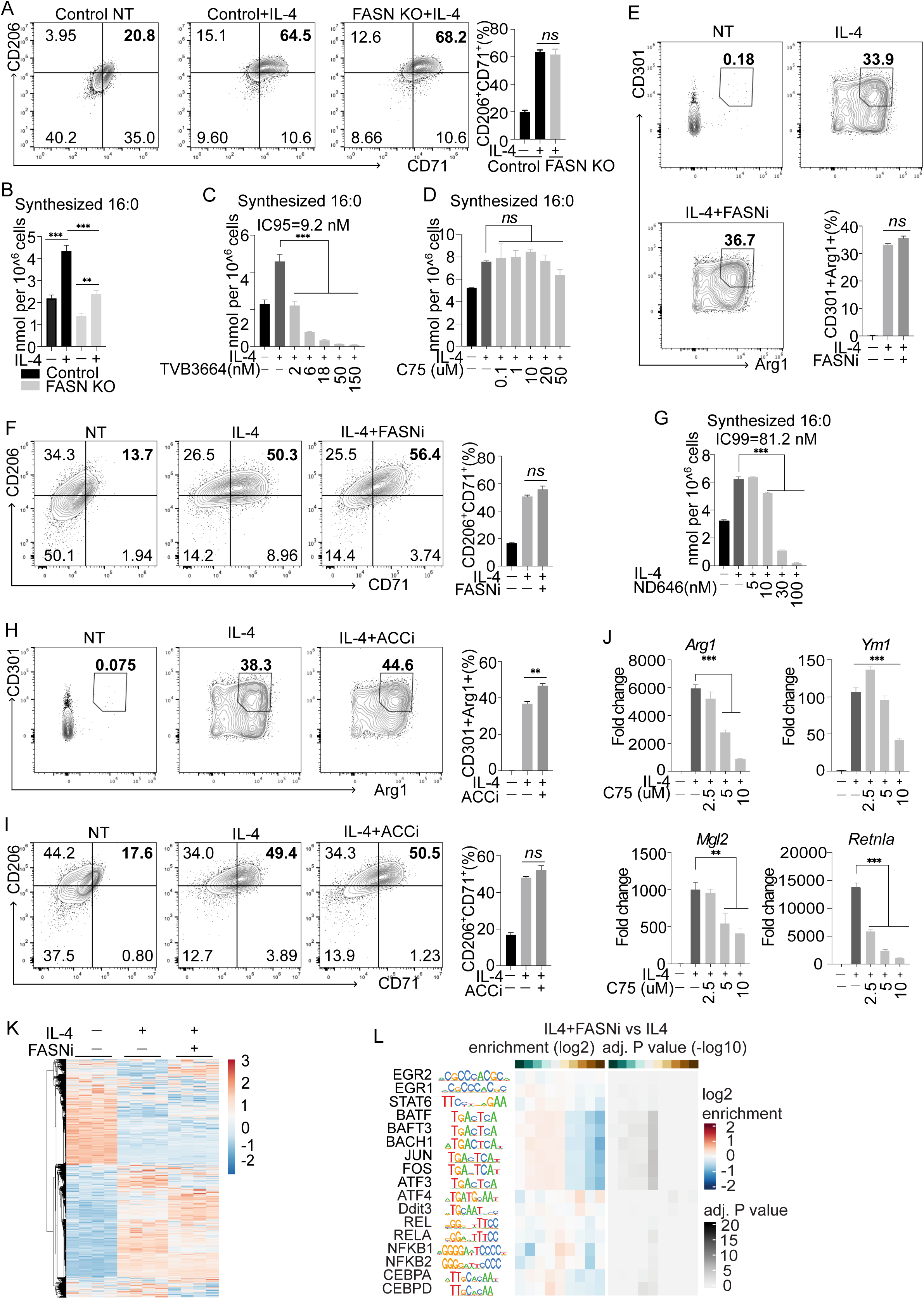
De novo lipogenic enzymes ACC and FASN do not perturb AAM polarization in vitro. A. Flow cytometry plots of LysMCre-control (indicated as Control) stimulated with or without IL-4 and LysMCre-Fasn^fl/fl^ BMDMs stimulated with IL-4 for 48 h. Bar graph (bottom right) depicts percent of double positive (CD206+/CD71+) BMDMs. B. Net synthesized palmitic acid (16:0) from LysMCre-control (WT) or LysMCre-Fasn^fl/fl^ BMDMs stimulated with or without IL-4 as indicated for 48 h. C. Net synthesized palmitic acid (16:0) from quiescent BMDMs (NT) or BMDMs stimulated with IL-4 ± indicated doses of FASN inhibitor TVB-3644 for 48 h. D. Net synthesized palmitic acid (16:0) from quiescent BMDMs (NT) or BMDMs stimulated with IL-4 ± indicated doses of FASN inhibitor C75 for 48 h. E. Flow cytometry plots of quiescent BMDMs (NT) and BMDMs stimulated with IL-4 ± FASNi as indicated for 48 h. Bar graph (right down) depicts percent of double positive (CDCD301+/Arg1+) BMDMs. F. Flow cytometry plots of quiescent BMDMs (NT) and BMDMs stimulated with IL-4 ± FASNi as indicated for 48 h. Bar graph (right) depicts percent of double positive (CDCD301+/CD71+) BMDMs. G. Net synthesized palmitic acid (16:0) from quiescent BMDMs (NT) or BMDMs stimulated with IL-4 ± indicated doses of ACC inhibitor ND646 for 48 h. H. Flow cytometry plots of quiescent BMDMs (NT) and BMDMs stimulated with IL-4 ± ACCi as indicated for 48 h. Bar graph (right) depicts percent of double positive (CDCD301+/Arg1+) BMDMs. I. Flow cytometry plots of quiescent BMDMs (NT) and BMDMs stimulated with IL-4 ± ACCi as indicated for 48 h. Bar graph (right) depicts percent of double positive (CDCD301+/CD71+) BMDMs. J. qPCR analysis of indicated AAM-associated genes from quiescent BMDMs (NT) or BMDMs stimulated with IL-4 ± indicated dosages of C75 for 48 h. K. Heatmap of RNA-seq data of quiescent BMDMs (NT) or BMDMs stimulated with IL-4 ± FASNi for 24 h. Genes (rows) were clustered according to correlation. L. Comparison of AAM- and SCDi-associated motifs in BMDMs stimulated with IL-4 ± FASNi for 24 h. Heatmaps representing enrichment (left) and adjusted p-values (right) for each of the sliding window bins (top). All experiments are from four biologic replicates per experimental condition. Heatmap scales are z-score for number of deviations away from the row mean. Rows are clustered using correlation distance and complete linkage. Gene expression and Flow cytometry studies are from three biologic replicates per experimental condition and are representative of greater than three experiments. Bar graphs are presented as mean ± SEM. *p < 0.05; **p < 0.01, ***p < 0.001.

**Figure S4.**
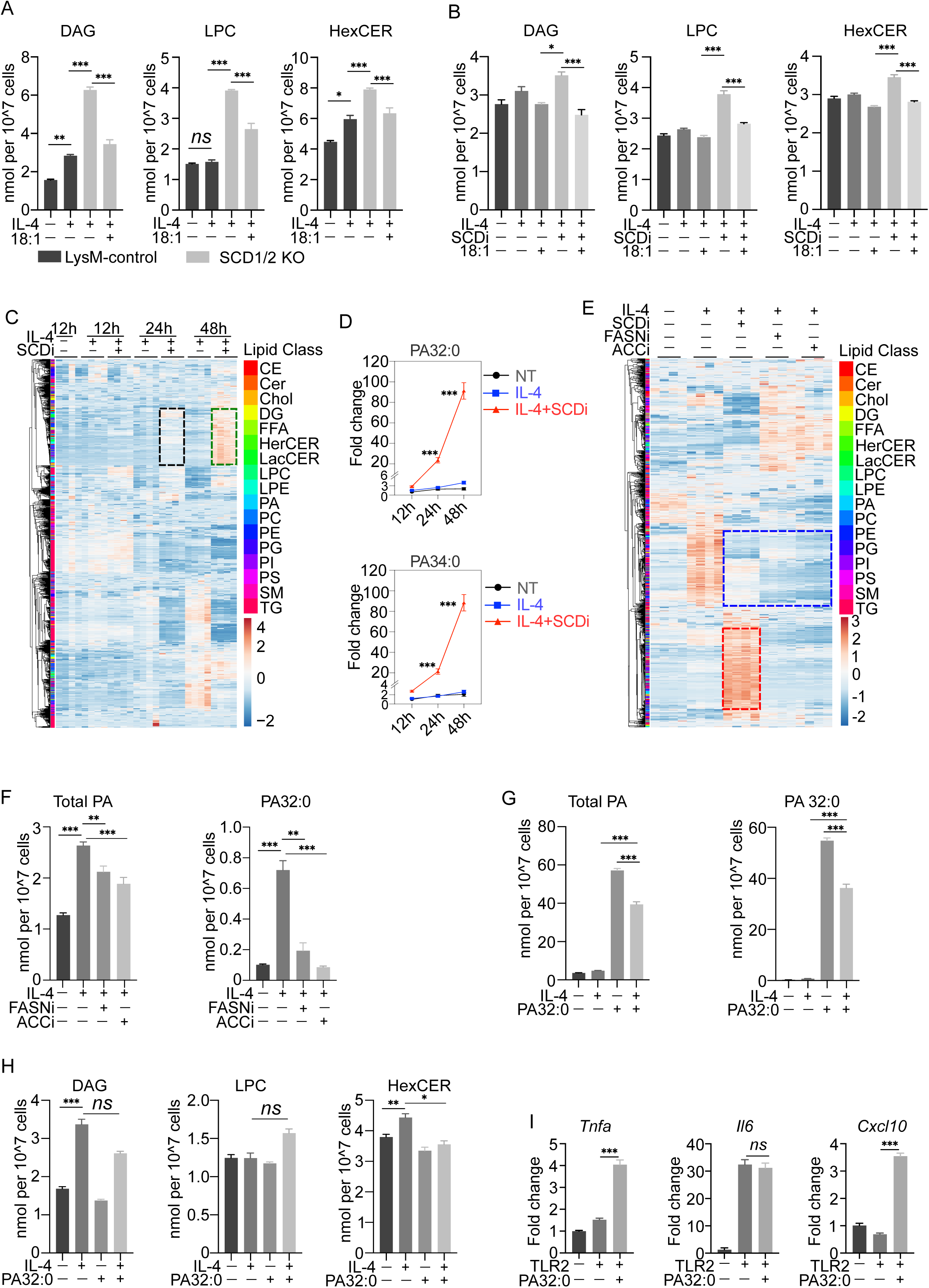
PA accumulation resulting from SCD loss impairs AAM polarization. A. Total diacylglycerol (DAG), lysophosphatidylcholine (LPC), and hexosylceramide (HexCER) measured by shotgun lipidomics from LysMCre control BMDMs stimulated with or without IL-4 and LysMCre-SCD1/2^fl/fl^ BMDMs stimulated with IL-4 ± 18:1 (25 μM) as indicated for 48 h. B. Total diacylglycerol (DAG), lysophosphatidylcholine (LPC), and hexosylceramide (HexCER) measured by shotgun lipidomics from from quiescent BMDMs (NT) or BMDMs stimulated with IL-4 ± SCDi or IL-4 ± SCDi with 18:1 (25 μM) as indicated for 48 h. C. Heat map of individual lipids quantified by shotgun lipidomics of quiescent BMDMs (NT) for 12h or BMDMs stimulated with IL-4 ± SCDi for 12h, 24h, 48 h. D. PA 32:0 (up) and PA 34:0 (down) measured by shotgun lipidomics quiescent BMDMs (NT) or BMDMs stimulated with IL-4 ± SCDi for 12h, 24h, 48 h. E. Heat map of individual lipids quantified by shotgun lipidomics from quiescent BMDMs (NT) or BMDMs stimulated with IL-4 ± SCDi or FASNi or ACCi as indicated for 48 h. F. Total phosphatidic acid (PA) (left) and PA 32:0 (right) measured by shotgun lipidomics from quiescent BMDMs (NT) or BMDMs stimulated with IL-4 ± FASNi or ACCi as indicated for 48 h. G. Total phosphatidic acid (PA) (left) and PA 32:0 (right) measured by shotgun lipidomics from quiescent BMDMs (NT) or BMDMs stimulated with IL-4 ± PA32:0 (10 μM) or PA 32:0 (10 μM) alone as indicated for 48 h. H. Total diacylglycerol (DAG), lysophosphatidylcholine (LPC), and hexosylceramide (HexCER) measured by shotgun lipidomics from quiescent BMDMs (NT) or BMDMs stimulated with IL-4 ± PA32:0 (10uM) or PA 32:0 (10 μM) alone as indicated for 48 h I. qPCR analysis of indicated pro-inflammatory genes of quiescent BMDM (NT) or BMDMs stimulated with TLR2 agonist ± PA 32:0 (10 μM) for 48 h. Lipidomics experiments are from four biologic replicates per experimental condition. Heatmap scales are z-score for number of deviations away from the row mean. Rows are clustered using correlation distance and complete linkage. Gene expression and flow cytometry studies are from three biologic replicates per experimental condition and are representative of greater than three experiments. Bar graphs are presented as mean ± SEM. *p < 0.05; **p < 0.01, ***p < 0.001.

**Figure S5.**
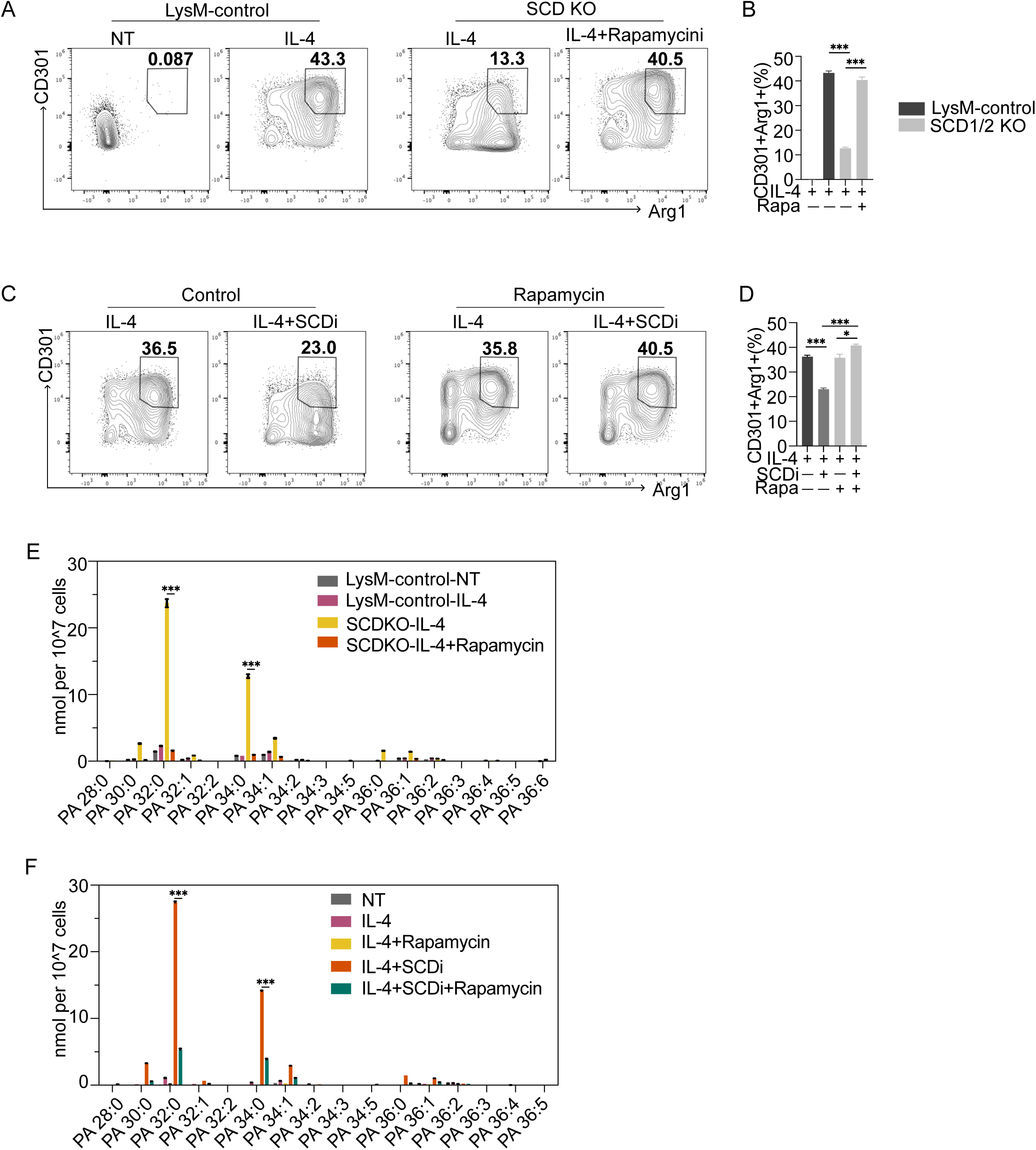
Inhibition of mTORC1 signal rescues AAM and PA homeostasis. A. Flow cytometry plots from LysMCre control BMDMs stimulated with or without IL-4 and LysMCre-SCD1/2^fl/fl^ BMDMs stimulated with IL-4 ± Rapamycin (10 nM) as indicated for 48 h. B. Bar graph depicts percent of double positive (CDCD301+/Arg1+) BMDMs as above. C. Flow cytometry plots from BMDMs stimulated IL-4 ± SCDi or IL-4 ± SCDi with Rapamycin (10 nM) as indicated for 48 h. D. Bar graph depicts percent of double positive (CDCD301+/ Arg1+) BMDMs as above. E. Amounts of individual PA species measured by shotgun lipidomics from LysMCre control BMDMs stimulated with or without IL-4 and LysMCre-SCD1/2^fl/fl^ BMDMs stimulated with IL-4 ± Rapamycin (10 nM) as indicated for 48 h. F. Amounts of individual PA species measured by shotgun lipidomics from quiescent BMDMs (NT) or BMDMs stimulated with IL-4 ± SCDi or IL-4 ± SCDi with Rapamycin (10 nM) as indicated for 48 h. Lipidomics experiments are from four biologic replicates per experimental condition. Heatmap scales are z-score for number of deviations away from the row mean. Rows are clustered using correlation distance and complete linkage. Gene expression and flow cytometry studies are from three biologic replicates per experimental condition and are representative of greater than three experiments. Bar graphs are presented as mean ± SEM. *p < 0.05; **p < 0.01, ***p < 0.001.

**Figure S6.**
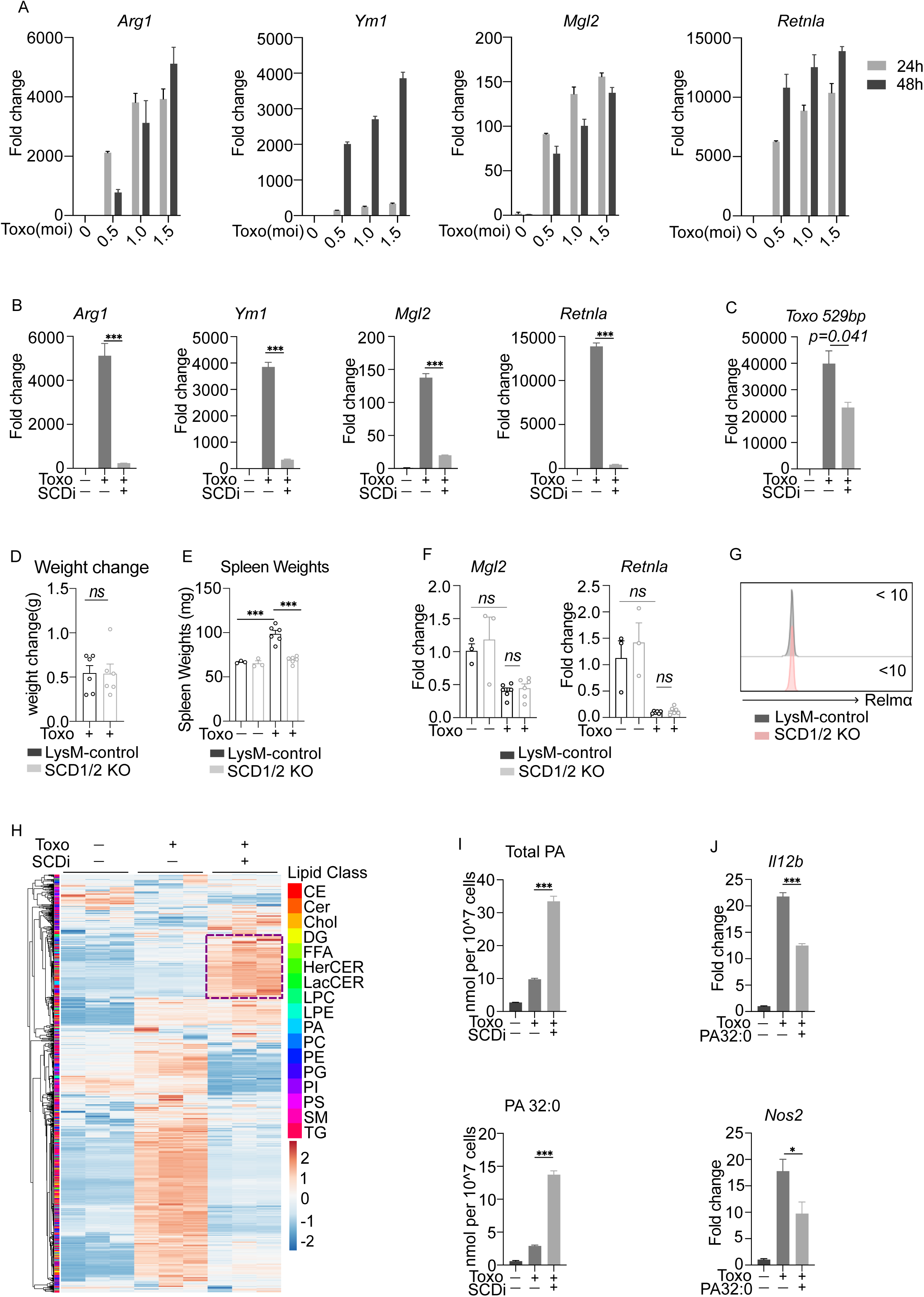
Loss of SCD disrupts AAM reprogram during Toxoplasma infection. A. qPCR analysis of select AAM-associated genes from quiescent BMDMs (no infection) or BMDMs infected with *Toxoplasma* at a MOI 0.5,1.0 and 1.5 for 24h and 48 h. B. qPCR analysis of select AAM-associated genes from quiescent BMDMs (no infection) or infected BMDMs (MOI=1.5) ± SCDi for 48 h. C. qPCR analysis of Toxoplasma gondii *529-bp repeat element* (*Toxo 529bp*) from quiescent BMDMs (no infection) or infected BMDMs (MOI=1.5) ± SCDi for 48 h. D. Bar graph depicts the weight change of LysMCre-control and LysMCre-Scd1/2^fl/fl^ mice infected with 1000 parasites for 5 days. E. Bar graph depicts the spleen weight of LysMCre-control and LysMCre-Scd1/2^fl/fl^ infected with 1000 parasites for 5 days. F. qPCR analysis of select AAM-associated genes from peritoneal cells collected by PBS lavage from LysMCre-control and LysMCre-Scd1/2^fl/fl^ mice infected with 1000 parasites for 5 days. G. Plot of MFI of CD206 and RELMα from peritoneal macrophages collected from LysMCre-control and LysMCre-Scd1/2^fl/fl^ mice infected with 1000 parasites for 5 days. H. Heat map of individual lipids quantified by shotgun lipidomics from quiescent BMDMs (no infection) or infected BMDMs (MOI=1.5) ± SCDi (10 nM) for 48 h. I. Total phosphatidic acid (PA) and PA 32:0 measured by shotgun lipidomics from quiescent BMDMs (no infection) or infected BMDMs (MOI=1.5) ± SCDi (10 nM) for 48 h. J. qPCR analysis of indicated pro-inflammatory genes from quiescent BMDMs (no infection) or infected BMDMs (MOI=1.5) ± PA 32:0 (10 μM) for 48 h. Gene expression and flow cytometry studies are from three biologic replicates per experimental condition and are representative of greater than three experiments. In vivo experiments are representative of n=6 mice per group repeated at least two times. Bar graphs are presented as mean ± SEM. *p < 0.05; **p < 0.01, ***p < 0.001.

## Notes

### Competing Interest Statement

The authors have declared no competing interest.

